# Prediction of Piconewton Receptor Tension Images using Deep Learning

**DOI:** 10.64898/2025.12.19.695587

**Authors:** Kartikey Kansal, Monica Umesh, Myrna Chang, Dominique Smith, Hemakshi Mishra, Juan Caicedo, Joshua M Brockman

## Abstract

Piconewton (pN) receptor forces govern many biological processes, but measuring these forces remains challenging. Molecular tension probes (MTPs) provide a sensitive means to measure pN cellular forces via fluorescence microscopy; however, MTPs are challenging to use and only forces transmitted through the probes are reported, complicating their use in heterogenous environments. Here, we present Tension Deep Learning (TensionDL), which leverages convolutional neural networks and image-to-image translation to predict pN receptor tension maps from images of cell morphology and the force-transducing protein vinculin. We validate the accuracy of TensionDL at the subcellular and cellular scales, demonstrate model accuracy across different substrate stiffnesses and cell types, and leverage TensionDL to make semi-quantitative predictions of cell mechanical output. Finally, TensionDL enables long-term mapping of pN receptor tension and infers tension distributions in heterogeneous environments in which some forces are not transduced through MTPs.

## Introduction

Cells sense and transmit piconewton (pN) scale forces via mechanically active receptors. These cellular forces regulate critical biological processes including coagulation^1^, cell differentiation^2^, and T cell antigen recognition^3,4^. We and others have generated molecular tension probes (MTPs) that transduce pN forces into fluorescence, enabling measurement of pN receptor tension magnitude and distribution^5–9^. In particular, DNA-based MTPs are sensitive and quantitative tension probes which utilize DNA hairpins flanked by a fluorophore-quencher pair^7^. When forces on the DNA MTP exceed the probe F_1/2_, defined as the equilibrium force that leads to a 50% chance of hairpin unfolding, the hairpin ruptures, separating the fluorophore from the quencher and producing a “turn-on” increase in fluorescence.

MTPs have limitations which hinder their use. First, they can be difficult to synthesize and are typically made at the nanomole scale hindering high-throughput applications and widespread adoption. Second, they suffer from imaging challenges including near single molecule fluorescence, rapid photobleaching, and force induced probe rupture at short time scales^7^. Most importantly, MTP measurements only report pN forces transmitted through the probe, limiting their use on physiological substrates. If MTPs were employed on substrates with native cell adhesion ligands, such as natural extracellular matrix (ECM), any forces transmitted to the ECM would not be reported. This is a critical limitation since mechanotransduction mechanisms depend on many properties of native ECM^10^, and disease-relevant processes like oncogenic transformation depend on physiological ECM mechanics^11^.

Recent advances in machine learning (ML) have enabled image-to-image translation, turning an input into a corresponding output: for example, image-to-image translation can translate a picture taken during the day into a corresponding picture taken at night^12^. Previous works have used image-to-image translation in biological applications such as generating histology^13,14^ or fluorescence images^15^ from unlabeled input images. Convolutional neural networks (CNNs) can predict fluorescent markers for nuclei, organelles, and even cell cycle state from transmitted-light images^15, 16^. This approach has recently been used to predict cellular forces from Traction Force Microscopy (TFM) maps and paired images of force transducing proteins^17^. However, TFM provides micrometer spatial resolution with nanonewton (nN) force sensitivity and thus cannot resolve receptor-level forces^18^. Additionally, TFM relies on homogenous, isotropic and known substrate properties which will ultimately limit its use in native environments. The combination of ML and MTPs offers an alternative solution.

We propose Tension Deep Learning (TensionDL) as a method to predict pN receptor forces from images of mechanically active proteins. Our core hypothesis is that cell morphology and the spatial distribution of mechanotransducing proteins encode sufficient information to reconstruct MTP fluorescence images. To train TensionDL, we produced paired images of cell morphology and the mechanotransducing protein vinculin as inputs, along with the desired output, MTP fluorescence images. We then modelled the problem as an image-to-image translation task to predict high-fidelity integrin-generated pN tension maps without the use of MTPs or knowledge of substrate mechanics. Here we validate the accuracy of TensionDL at the cellular and subcellular scales on both glass and soft hydrogels matching tissue stiffness. We employ this capability to bypass limitations of MTPs by enabling multi-hour timelapse imaging of cellular mechanics and by inferring integrin forces transmitted to ECM coated substrates in which most forces are transmitted to the ECM and thus are not observed.

## Results

### TensionDL predicts the distribution of MTP fluorescence signal

We synthesized previously reported^7^ 4.7pN F_1/2_ DNA MTPs presenting the integrin adhesion ligand cyclo-Arg-Gly-Asp (cRGD) and labeled with Cy3B-BHQ2 a fluorophore-quencher pair (**Fig. 1a**, **Supplementary Fig. 1, Supplementary Table 1**). The F_1/2_ is defined as the equilibrium force that leads to a 50% chance of probe opening. We seeded mouse embryonic fibroblast cells expressing GFP-vinculin (MEF GFP-vin) on substrates coated with the MTPs and allowed them to spread for 15 min before imaging (**Supplementary Fig. 2**). As model inputs, we acquired images of cell-surface contact area via reflection interference contrast microscopy (RICM) and the location of Vinculin via total internal reflection fluorescence (TIRF) microscopy. We also acquired the corresponding paired image of the desired output, Cy3B MTP fluorescence, which served as our ground truth (GT) for model training (**Fig. 1b**). We selected 1312, 256 x 256 pixel regions of interest (ROIs) from our data set (**Supplementary Fig. 3, 4**) which were allocated to training (797 ROIs), validation (143 ROIs), and test (371 ROIs) datasets. The training and validation data sets were supplemented with 30 and 34 images of cell-free background, respectively. Employing the previously reported model framework CytoDL from the Allen Institute (https://github.com/AllenCellModeling/cyto-dl), we trained a U-Net convolutional neural network to perform an image-to-image translation task which translates RICM and GFP-vin images into maps of 4.7pN MTP fluorescence signal, inferring the distribution of integrin generated forces (**Fig. 1b, Supplementary Fig. 5**). This model, which we named Tension Deep Learning (TensionDL), successfully predicts images of cells from the test dataset which bear a striking resemblance to the GT 4.7pN tension images (**Fig. 1c**).

**Figure 1:**
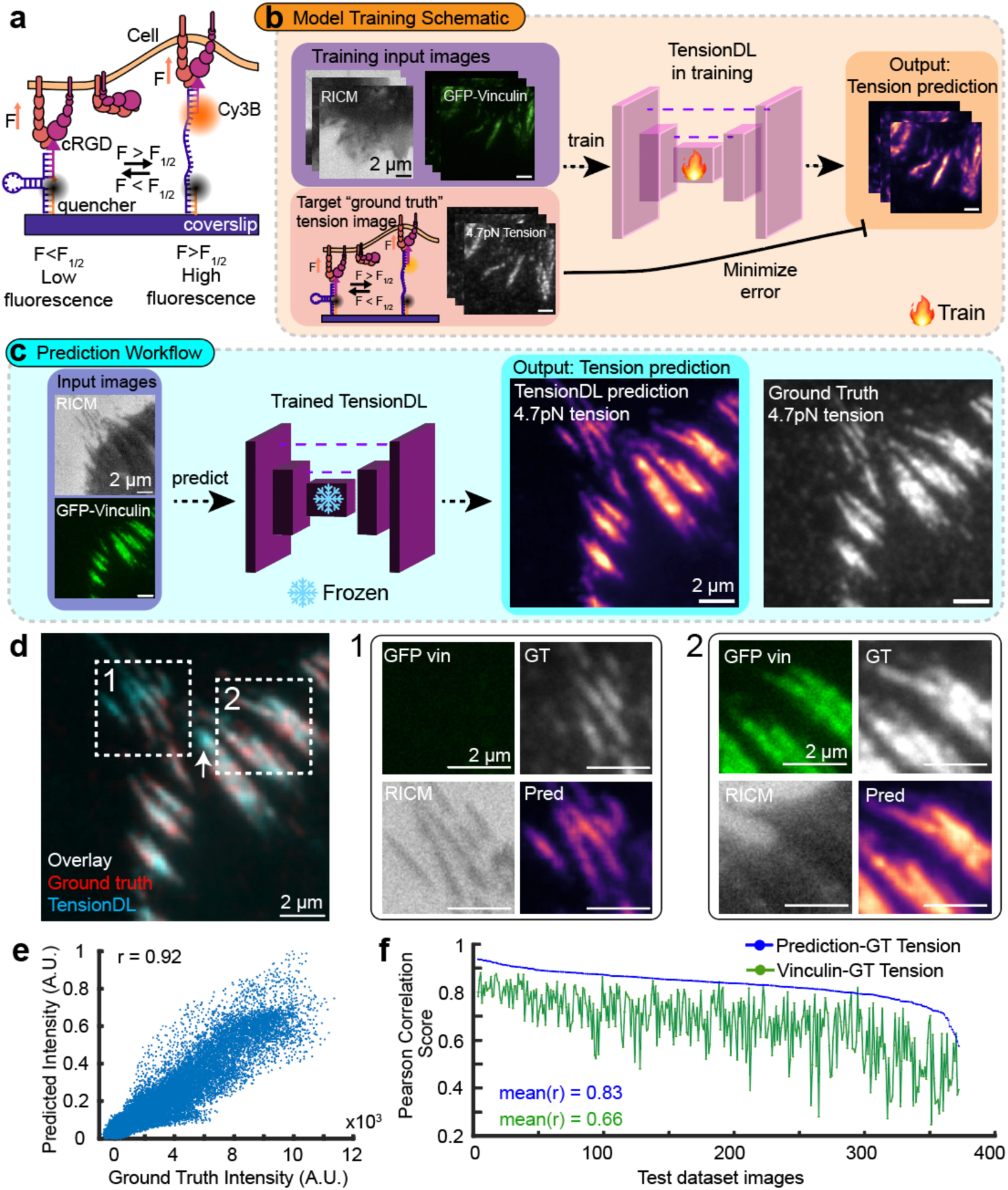
TensionDL predicts the distribution of piconewton receptor forces. **a**, Schematic of the DNA MTPs which report pN forces by fluorescence when force is exceeding the F_1/2_ open the probe. Probes with F_1/2_ = 4.7pN were used to create the training data set. **b**, Schematic of TensionDL model training, Representative images of MEF surface contact area (RICM), GFP-Vinculin fluorescence and 4.7pN MTP fluorescence **c,** TensionDL prediction (magma colormap) and ground truth image (grayscale) for a cell withheld from the model during training. **d**, Red-cyan overlay of Ground Truth and Prediction for cell in (c) (normalized to their max values), respectively, where white indicates perfect overlay. Zoomed ROIs 1 and 2 are indicated by dashed boxes. **e**, Scatterplot of ground truth versus predicted intensities for the cell shown in (**c**) and (**d**). The pixelwise pearson correlation for this cell is r=0.92. **f**, Plot of cell ID (x-axis) versus pixelwise pearson correlation (y-axis) for ground truth versus prediction (blue) and ground truth versus vinculin-GFP (green) for all 371 cells in the test dataset. Data in (**c-e**) are representative samples of the cells in the test dataset, sourced from *n*=6 independent experiments.

We next sought to determine the accuracy of the TensionDL predictions. Overlaying GT (red) with the prediction (blue) reveals striking agreement between the images (white, **Fig. 1d**). Interestingly, TensionDL predicts tension even in areas where GFP-vin signal is not apparent (**Fig. 1d**, ROI 1, **Supplementary Fig. 6**). A pixelwise scatterplot of GT images versus TensionDL prediction produces a positive correlation (**Fig. 1e, r** = 0.92 for cell shown). We extended this analysis to all 371 cells (*n* = 6 independent experiments) in the test dataset that were withheld from the TensionDL model during training, revealing an average pixelwise Pearson correlation of r = 0.83 between GT 4.7pN tension and TensionDL prediction (blue line, **Fig. 1f**). For comparison, we computed Pearson correlations between GFP-vin fluorescence and measured GT 4.7pN tension signal, revealing an average Pearson of 0.66 (green line, **Fig. 1f**). TensionDL predictions consistently outperform GFP-vin as a predictor of tension location (**Fig. 1f**).

### Prediction of pN forces at cellular and subcellular scale

To rigorously evaluate the predictive performance of TensionDL, we compared its output to ground-truth (GT) receptor tension maps at both the cellular and subcellular levels. For quantitative evaluation, both GT and predicted images were segmented into discrete “tension zones” (TZs), defined as 8-connected high-intensity regions exceeding 60 pixels in size (**Fig. 2a, b**). GT segmentation masks (color-coded) and segmented predicted zone (white outlines) reveal significant correspondence between experimentally measured and computationally inferred force domains (**Fig. 2b, c**). We quantified this overlap using the Jaccard similarity index^19^, which measures the intersection-over-union between paired zones. In the representative cell shown, TensionDL correctly predicted 14 of 16 zones, yielding an average Jaccard index of 0.58 (**Fig. 2c**), falling within a range typical for high-performing deep-learning models in biological image segmentation^20^. Across the full 371 image test dataset, the model correctly identified 5,527 of 7,531 GT zones (73.4%), while 26.6% were either missed or over-segmented (**Fig. 2d**). Notably, many of the missed or mismatched zones primarily correspond to small low intensity GT features (**Fig. 2b**, red zones). The centroid displacement between corresponding GT and predicted zones remained small (mean ∼ 0.2 µm; **Fig. 2f**) further indicating good agreement at the subcellular scale. Additionally, the area of Predicted and GT zone are strongly correlated (r = 0.91) (**Fig. 2e**).

**Figure 2:**
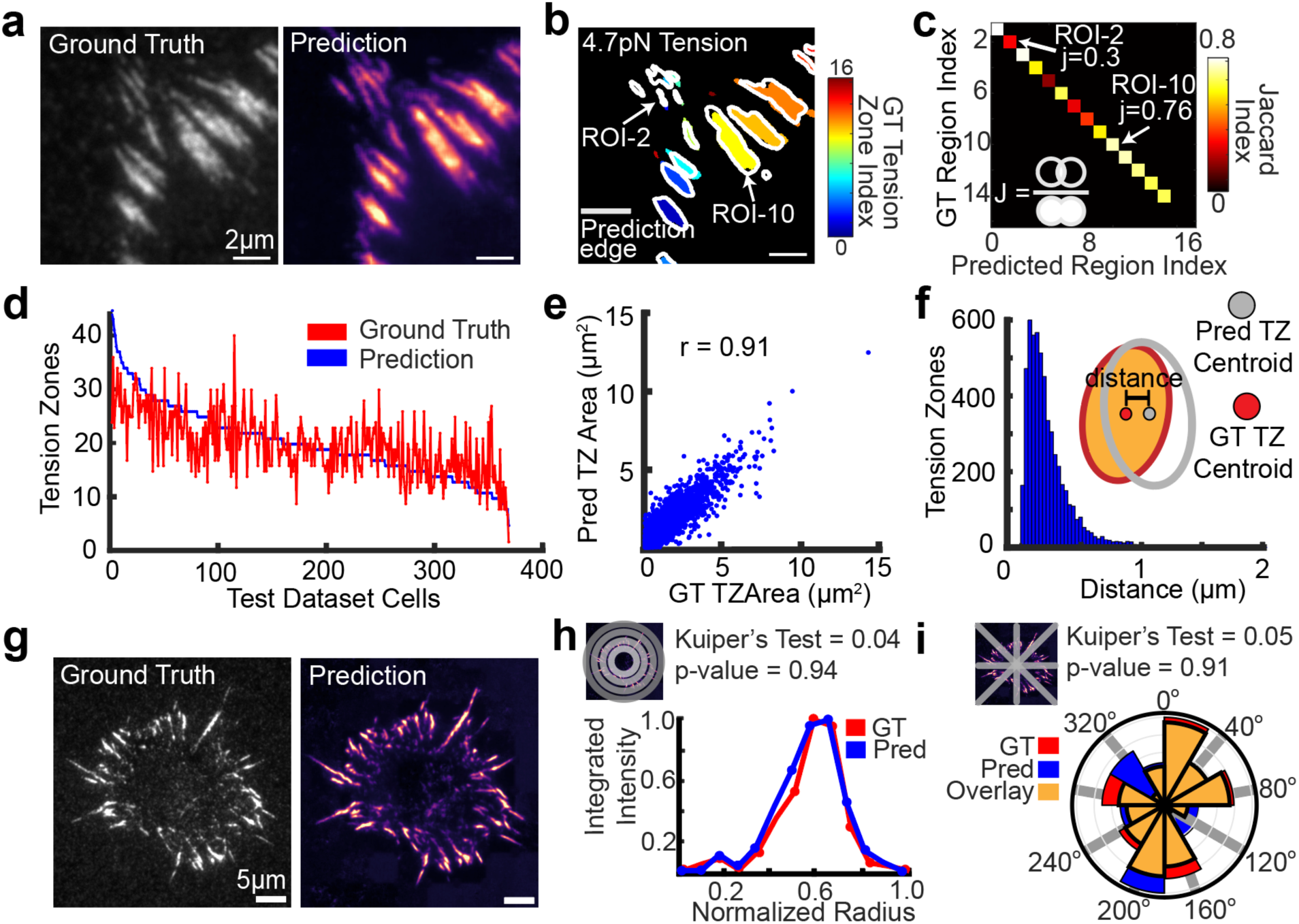
TensionDL predictions are accurate at the cellular and subcellular scales. **a**, Ground Truth (left), TensionDL prediction (right). **b,** Segmented tension zones of cell shown in **(a).** ROI label for the ground truth is indicated by color while edges of predicted tension zones are shown by gray edges. **c**, Heatmap of Jaccard Index for tension zones identified in (**b**). ROIs with high jaccard index (ROI-10) and low jaccard index (ROI-3) are indicated in both (**b**) and (**c**). **d**, Plot of number of tension zones identified (y-axis) for ground truth (red) and TensionDL prediction (blue) versus cell index (x-axis) for 371 cells in the test dataset. **e**, Scatterplot of masked ground truth versus prediction tension zone area for 7531 tension zones. **f**, Histogram of intensity weighted centroid distance between TensionDL and ground truth for all 7531 tension zones. **g**, Representative whole cell Ground Truth (left) and TensionDL prediction (right). **h**, Integrated normalized intensity for radial bins of ground truth (red) and prediction (blue), and (**i**) Angular histogram plot of distribution of pN receptor for the cell shown in (**g**). In (**i**), yellow represents overlay. Note that the cell in (**a**) and (**b**) also depicted in Fig 1c. (**a**-**f**) are representative of 371 cells in the test dataset (*n*=6 independent experiments). (**g**-**i**) are representative of 95 whole cell TensionDL predictions sourced from *n*=6 independent experiments.

At the whole-cell scale, TensionDL accurately captures global force organization. For 95 full-cell maps, we computed the radial and angular distributions of receptor tension signal (Fig. 2g). The radial intensity profile of the prediction closely overlapped with that of the GT map (Kuiper’s test p = 0.94; **Fig. 2h**), indicating that the model faithfully reproduces the overall distribution of tension from the cell edge toward the center. Likewise, the angular distribution of predicted tension (**Fig. 2i**) preserved the polarized distribution of the GT pattern (Kuiper’s test p = 0.91). Collectively, these results demonstrate that TensionDL achieves robust, multi-scale accuracy in predicting receptor tension distributions.

### TensionDL generalizes to new contexts

We next evaluated whether TensionDL could generalize to experimental conditions beyond those present in its training dataset. The original model was trained exclusively on data collected from glass substrates (Young’s modulus ≈ 60 GPa), which are many orders of magnitude stiffer than biological tissues (0.1–100 kPa)^21^. To test model transferability to physiologically relevant stiffnesses, we fabricated molecular tension probe (MTP) functionalized polyacrylamide (PAA) hydrogels with controlled elastic moduli of 21.9, 35.2, and 53.3 kPa (**Fig. 3a, Supplementary Table 4**). These gels were functionalized with cRGD-linked hairpin probes following an adaptation of established surface-coupling and hybridization protocols^22^. Because the optical path now runs through the hydrogel, we were unable to perform RICM imaging, and thus retrained a variant of TensionDL using only vinculin fluorescence as input. For reference, **Supplementary Table 3** details which model version was used for each figure. TensionDL maintained strong predictive performance across all stiffnesses (**Fig. 3b–d**). Predicted maps exhibit good agreement with ground truth measurements with pixelwise Pearson correlation between predicted and GT maps of r = 0.73 for 21.9 kPa, r = 0.82 for 35.2 kPa, and r = 0.87 for 53.3 kPa. As expected, predictive fidelity decreased on low stiffness gels which differ more significantly from the conditions used during training.

**Figure 3:**
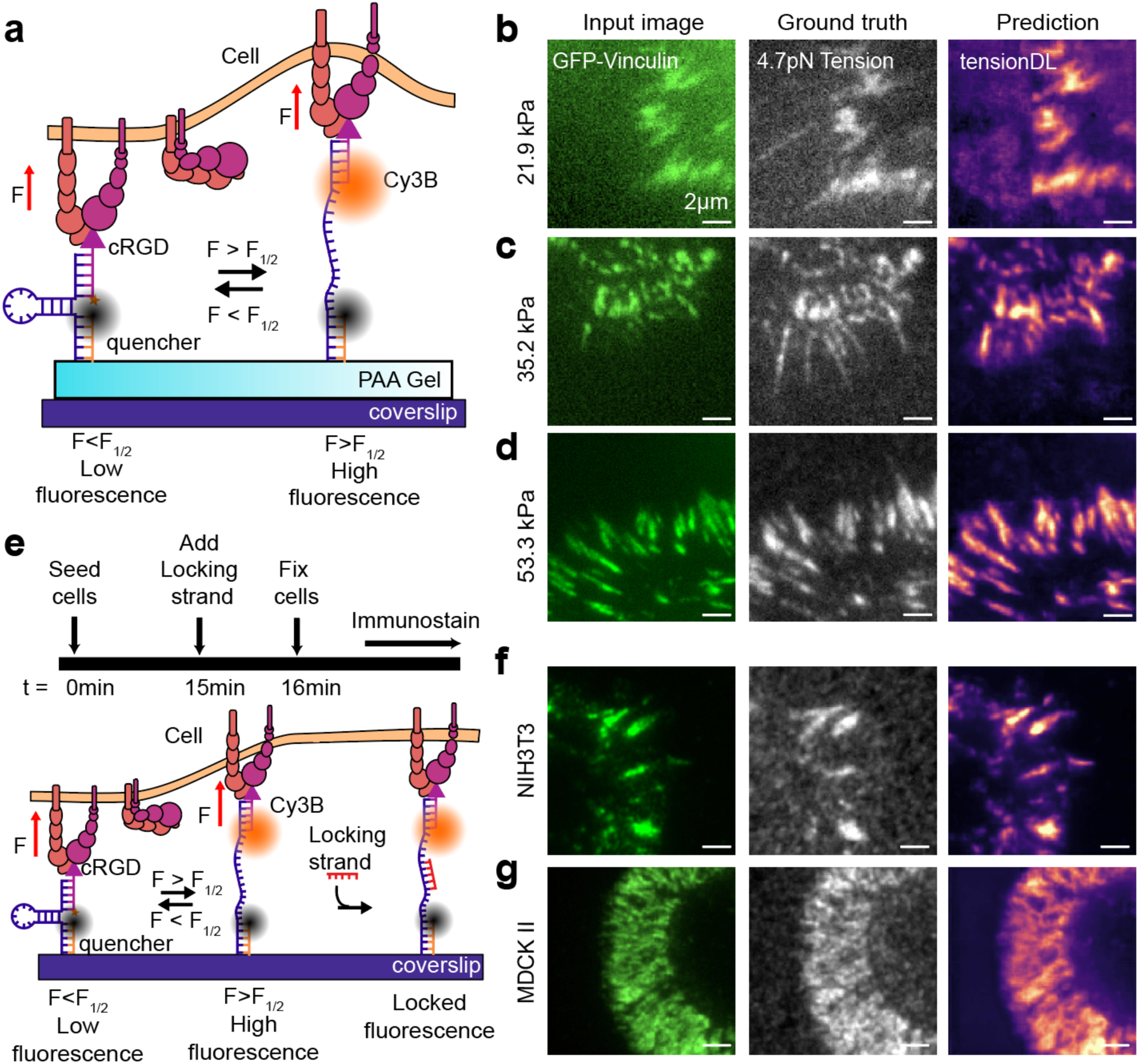
TensionDL generalizes to soft hydrogels, immunofluorescence detection of focal adhesion proteins, and new cell types. **a**, Schematic of DNA MTP on a soft PAA hydrogel. From left to right: Input GFP-Vinculin image, measured MTP fluorescence, and TensionDL prediction for MEF GFP-Vinculin fibroblasts seeded on (b) 21.9kPa, (c) 35.2kPa and (d) 53.3kPa PAA hydrogels conjugated with DNA MTPs, **e.** Schematic of DNA MTP locking and immunofluorescence detection of Vinculin. From left to right: Alexa488 anti-Vinculin stain, GT MTP Tension signal, and TensionDL prediction for (**f**) NIH3T3 fibroblasts and (**g**), MDCKII cells, respectively. Data shown are representative of **(b)** 65 cells (*n*=3 independent experiments), **(c)** 72 cells (*n*=3 independent experiments), **(d)** 97 cells (*n*=3 independent experiments), **(f)** 70 cells (*n*=3 independent experiments), **(g)** 65 cells (*n*=3 independent experiments).

To further assess model generalizability, we next tested TensionDL against different cell types in which Vinculin was detected using immunofluorescence. We seeded NIH 3T3 fibroblasts and MDCK II epithelial cells on cRGD-functionalized MTP substrates, fixed the cells, and stained for vinculin. To preserve tension during permeabilization and staining, we added a locking strand^23^ for 1 minute prior to fixation, preventing open hairpins from refolding during staining. We used the resultant Vinculin immunostaining data as an input to TensionDL yielding high quality MTP maps (**Fig. 3f, g**). TensionDL predictions for both NIH 3T3 (**Fig. 3f**) and MDCK II (**Fig. 3g**) cells closely resemble the GT maps. We observed strong pixelwise correlations between predicted and GT signals (r = 0.78 for NIH 3T3; r = 0.90 for MDCK II), suggesting that the model generalizes across different cell types. Importantly, TensionDL’s compatibility with immunofluorescence may be important in situations where genetic manipulation of cells is not feasible. Together, these findings indicate that TensionDL is robust to substantial shifts in imaging modality, substrate stiffness, and cellular phenotype.

### TensionDL enables mechanical imaging on ECM coated substrates

We next sought to use TensionDL to circumvent common limitations of MTPs. MTP imaging is complicated by photobleaching, and probe rupture (**Supplementary Fig. 7, 8**), underscoring the challenge of direct long-term mechanical imaging with tension probes. Another persistent limitation of MTPs is that they only report the subset of forces transmitted through the probe, limiting their use in complex environments where native extracellular matrix (ECM) ligands compete for receptor binding. To overcome these restrictions, we evaluated whether TensionDL could enable long duration measurements which infer the distribution of pN receptor forces on surfaces containing both synthetic MTPs and natural ECM proteins. We adsorbed fibronectin, an RGD-containing ECM protein, onto MTP-functionalized coverslips^24^ (**Fig. 4a**). On these hybrid substrates, fibronectin and MTPs compete for integrin engagement and forces transmitted through fibronectin are not observable by MTPs. As expected, increasing fibronectin concentration reduced MTP signal intensity (**Supplementary Fig. 9**), indicating competition between MTPs and fibronectin. We sought to determine whether the fibronectin perturbed the inputs to TensionDL. Accordingly, we extracted 384 high-dimensional features from the input channels (vinculin and RICM) using a pretrained DINOv2 transformer as an unbiased quantification of cellular morphology. We reasoned that if fibronectin does not perturb model input, then TensionDL predictions remain valid even when we cannot directly observe all cellular forces. We visualized these extracted features using a UMAP projection (**Supplementary Fig. 9**). UMAPs projections generated from features extracted from the vinculin input channel (**Supplementary Fig. 9d**) did not exhibit clustering in response to fibronectin treatment, indicating that model input features were relatively unaffected. We therefore concluded the TensionDL predictions should remain valid under these conditions. Importantly, to the extent that we could still measure MTP fluorescence despite competition with fibronectin and photobleaching, TensionDL predictions continued to correlate positively with MTP measurements (**Supplementary Fig. 10**).

**Figure 4:**
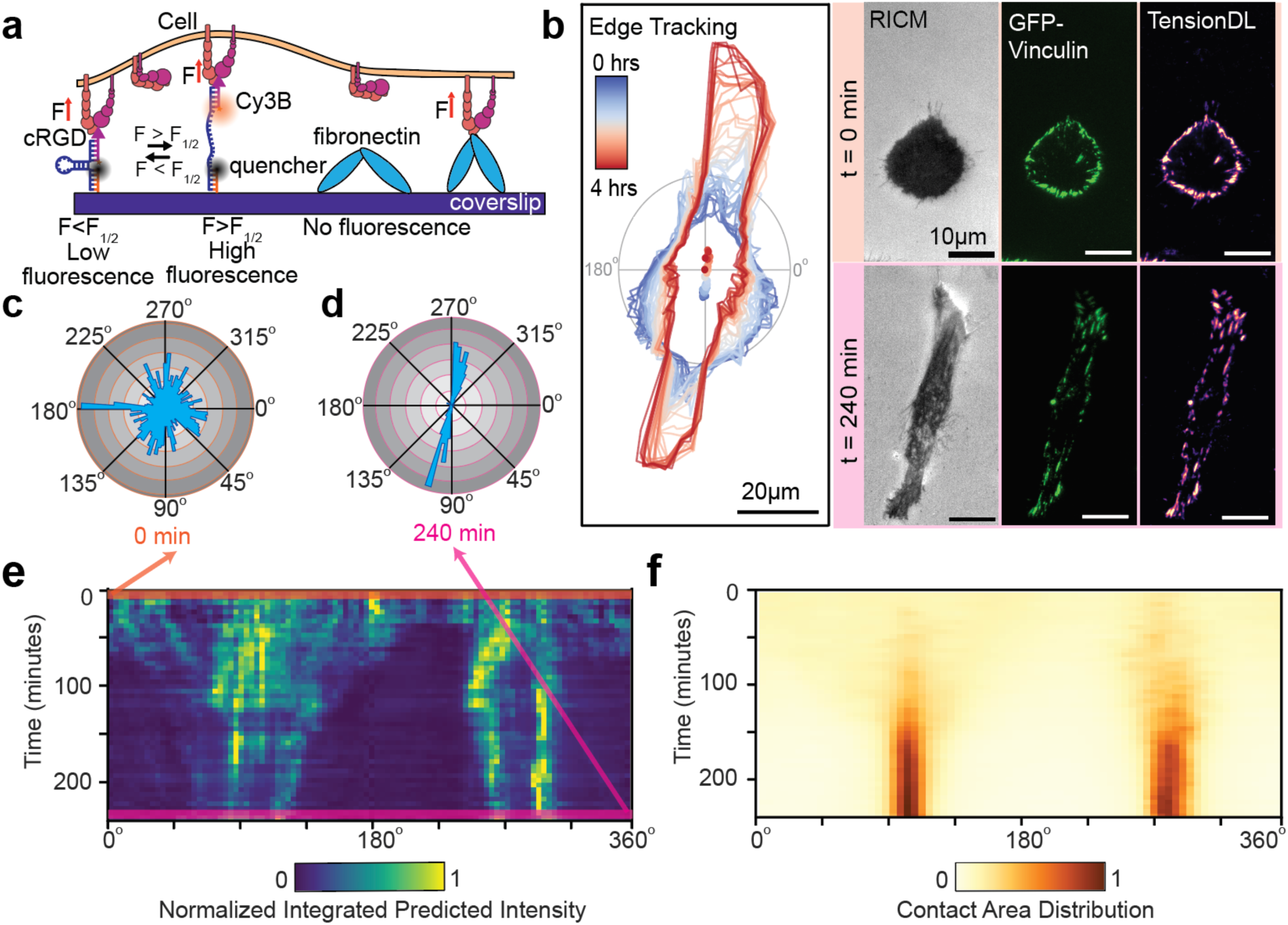
TensionDL long term imaging on heterogeneous substrates coated with ECM protein. **a.** Schematic of coverslip coated with DNA-MTP and the ECM protein fibronectin**, b.** Left: edge tracking of MEF GFP-Vinculin cell where color indicates the time of observation, right: RICM, GFP–vinculin, and TensionDL predictions at (top right) t = 0 and (bottom right) 240 min, **c.** Angular histogram of intensity weighted TensionDL prediction at 0 min and **(d)** 240 min, **e.** Angular kymograph heatmap of normalized integrated intensity of TensionDL predictions for cell in (**b)**, binned by angle (x-axis) and tracked over 4 h (y-axis), **f.** Angular kymograph heatmap of RICM-based contact distribution for the same cell over 4 h.

Finally, we employed TensionDL to predict force maps every 5 minutes over a 4-hour imaging period on a heterogeneous surface coated in both fibronectin and tension probes (**Fig. 4a, b, Supplementary Video 1**). To visualize the evolution of tension over time, we divided the cell into 2° angular wedges and computed the integrated intensity within each wedge. Angular histograms of the distribution of predicted forces are provided for 0 min (**Fig. 4c**) and 240 min (**Fig. 4d**). A similar calculation was performed on cell surface contact area using the RICM image. To visualize dynamics over the 4-hour imaging window, we generated an angular kymograph for TensionDL predictions (**Fig. 4e**) and cell surface contact area (**Fig. 4f**). The radial histogram for 0 min and 240 min is the top and bottom line in this kymograph, respectively (**Fig. 4e**). Strikingly, TensionDL revealed the early emergence of mechanical polarity, with forces aligning along the 90°/270° axis within the first 15 minutes of imaging (**Fig. 4e**). This axis persisted for the entire 4-hour imaging window. Cell polarization along this axis was not evident until approximately 100 minutes into the timelapse. This long-term measurement of forces on a heterogeneous surface is not possible with standard DNA MTPs.

### TensionDL enables relative cell to cell tension comparison

Finally, we sought to determine whether TensionDL could enable relative mechanical comparisons, e.g. comparing integrated tension signal between two cells. To our surprise, although our model captured the spatial distribution of forces accurately, it wasn’t as accurate in cell-cell comparison. Standard image-to-image translation models do not focus on quantitative comparisons, because their training objectives are strictly per-pixel reconstruction losses^25^. We identified two confounders in our data: 1) input and target data normalization operations common in image-to-image models obscured quantitative intensity information, 2) technical variation in GT MTP images led to biases in intensity information.

To mitigate these confounders, we employed a local and global normalization across our entire MTP training dataset. The local normalizations consisted of dividing each image by the intensity of the cell-free quenched MTP intensity for that image, which accounts for technical variation in the experiment. The global normalizations, which followed the local normalizations, subtracted the median and divided by the maximum pixel value across the training dataset. The effect of this operation was to align the intensities between different cells. We trained a model on this normalized dataset, which we refer to as TensionDL-Q (TensionDL quantification). To test TensionDL-Q, we generated “low” MTP intensity data using moderate pharmacological perturbations that alter, but do not eliminate, cellular forces (**Fig. 5**). MEF GFP–vinculin (GFP–vin) cells were treated with either Latrunculin B (LatB), which sequesters G-actin and prevents filament assembly, or CK666, an inhibitor of Arp2/3-mediated branched actin polymerization (**Fig. 5a–f**). Following LatB treatment, GT tension intensity dropped sharply (t = 5 min; **Fig. 5a**, green dashed line). We computed whole cell integrated tension for both GT and TensionDL-Q predictions as a metric of cell mechanical output. TensionDL-Q predicts a decrease in tension upon drug treatment (**Fig. 5a, b**). At the population level, TensionDL-Q predicts higher integrated intensities before LatB treatment than it does after treatment, although it underestimates the magnitude of the decrease (**Fig. 5c**). This substantiates the ability of TensionDL-Q to perform relative comparison in MTP intensity. CK666 treatment produced a slower and more gradual reduction in tension, consistent with dampened lamellipodial activity and loss of branched-actin support. TensionDL-Q reproduced the dynamics (**Fig. 5d, e**) and magnitude (**Fig 5f**) of the tension decrease following CK666 treatment. TensionDL-Q was unable to recapitulate the decrease in MTP signal following 5uM Y27632 RHO-kinase inhibition, although spatial maps of MTP intensity retained their fidelity (**Supplementary Fig. 12**). We hypothesize that this occurred because the model was trained exclusively on unperturbed cells.

**Figure 5:**
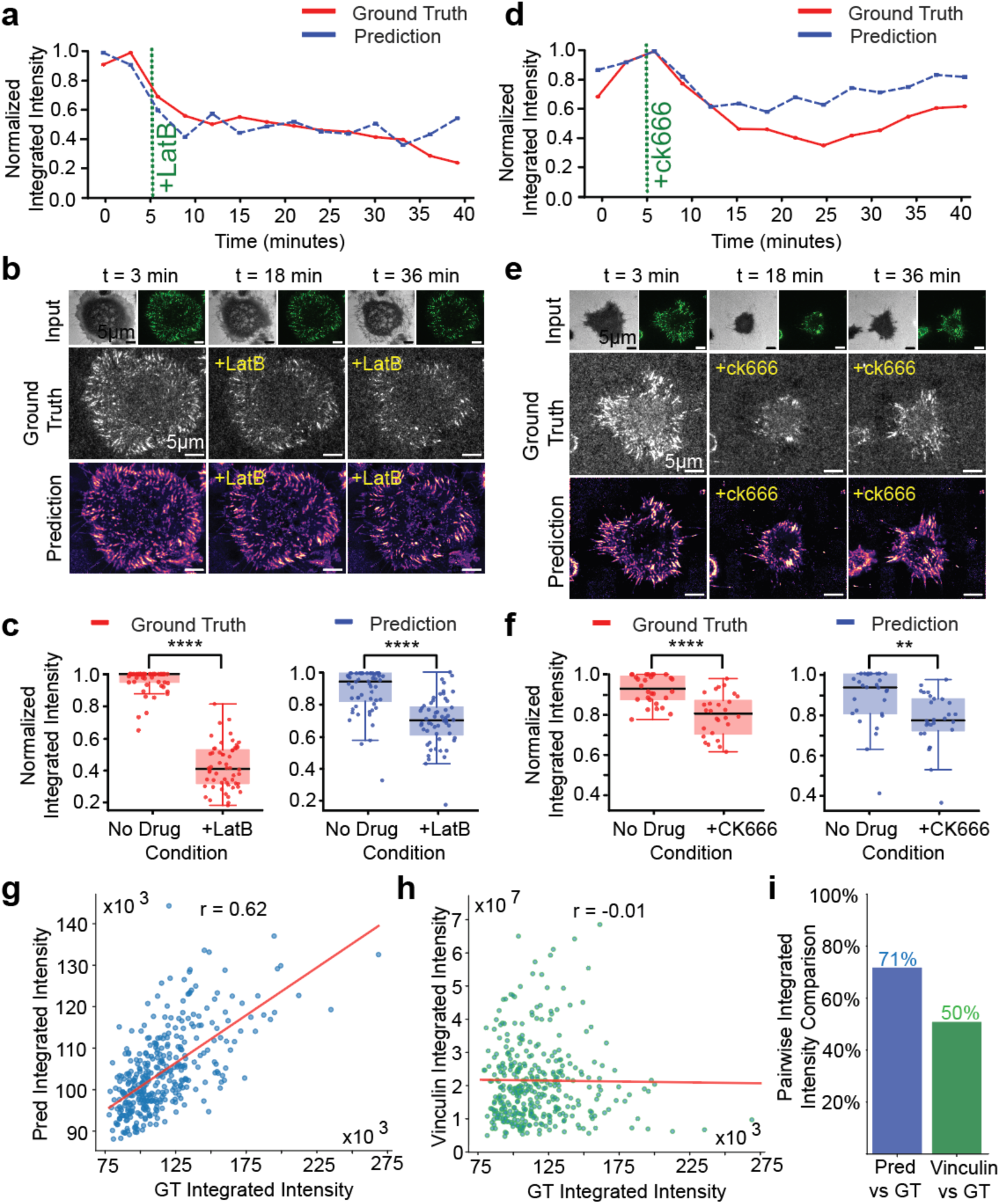
TensionDL predicts effects of pharmacological perturbation and relative cell to cell MTP intensity. **a, d.** Timelapse traces of normalized integrated MTP-tension with drug addition at t = 3 min (green dashed line): LatB (1 µM) in **a** and CK666 (150 µM) in **d**. Red = GT MTP-tension; blue = TensionDL predictions. **b, e.** Representative images at t = 3, 18, and 36 min showing input (top), GT MTP-tension maps (middle), and TensionDL predictions (bottom) for LatB **(b)** and CK666 **(e). c**, **f.** Box-and-whisker plots of per cell normalized integrated MTP-tension before and after treating with respective cytoskeletal inhibitor for GT and TensionDL prediction across **(c)** *n* = 71 cells, 3 experiments; **(f)** n = 70 cells, 3 experiments; significance was determined by Welch’s t-test (****P < 0.0001; **P < 0.01). **g**. Scatter of predicted (y-axis) vs GT integrated MTP-tension (x-axis) with least-squares fit; Pearson r=0.62. **h.** Scatter of GFP–vinculin integrated intensity (x-axis) vs GT MTP-tension (y-axis) with least-squares fit; Pearson r=-0.01. **i.** Bar plot of pairwise integrated intensity comparison agreement with GT for TensionDL predictions (blue) and vinculin (green); percentages labeled.

We next evaluated the ability of TensionDL-Q to predict the magnitude of whole cell integrated MTP signal in unperturbed cells, which we commonly employ as a readout of cell mechanics. Across the full test set of 371 cells, TensionDL-Q prediction integrated intensities demonstrated a positive correlation with experimental measurements, achieving r = 0.62 between predicted and ground-truth (GT) integrated intensities (**Fig. 5g**). In contrast, the integrated intensity of vinculin is not a good reporter of MTP fluorescence (r = –0.01; **Fig. 5h**). To further assess the ability of TensionDL-Q to compare cell to cell MTP intensity, we performed a pairwise comparison analysis between every possible pair in our 371-cell test set. TensionDL-Q predicted the cell with higher MTP fluorescence 71% of the time, while vinculin intensity achieved only 50% accuracy, no better than a coin flip (**Fig. 5i**). Together, these data suggest TensionDL-Q enables relative cell to cell mechanical comparisons.

## Discussion

Here, we demonstrate that TensionDL, a CNN trained on inputs of cell morphology and the spatial distribution of mechanically active proteins, can infer spatial maps of pN receptor forces. We validated the accuracy of TensionDL predictions at the cellular and subcellular scale by comparison with MTP measurements that were withheld from the model during training. TensionDL generalized well across different substrate stiffnesses and cell types. Additionally, the imaging modalities required for TensionDL such as RICM and fluorescence imaging of protein locations are broadly accessible to the biological field. We anticipate that TensionDL will democratize access to piconewton receptor force measurements. Additionally, TensionDL may enable mechano-imaging in high-throughput applications, allow imaging of cellular dynamics at high frame rates, and will permit long term imaging of cellular forces.

TensionDL circumvents many of the limitations of existing techniques to measure cellular forces. MTPs measure receptor-ligand forces with piconewton sensitivity, but suffer from photobleaching, and only report forces transmitted through the probe–receptor junction^7,9,26^. Genetically encoded FRET-based tension sensors^5,27^ also suffer from limited signal-to-noise ratio and the risk of perturbing protein function, constraining their use^28,29^. Traction force microscopy (TFM), quantitatively reconstructs traction stresses from substrate deformations, but typically requires the assumption of time invariant, isotropic, homogenous mechanical properties, hindering use in complex environments^30–33^. TensionDL overcomes many of the above limitations. Instead of tracking probe opening or substrate deformation, the network integrates cues such as focal-adhesion architecture and cell shape to recover the mechanical output of the cell. As a result, TensionDL functions without knowledge of substrate mechanical properties and is not susceptible to photobleaching of MTPs during imaging.

TensionDL comes with some limitations. The current TensionDL model was trained on MEF GFP-Vinculin fibroblasts and generalized well to predict forces in 3T3 fibroblasts and MDCKII epithelial cells. Vinculin is commonly associated with focal adhesions and actomyosin contractility. It is likely that TensionDL will perform well with cell types that exert forces through similar mechanisms. In contrast, many cell types such as T cells leverage actin polymerization for force generation^3^, thus future work should explore training TensionDL variants to predict forces using actin images or images of other mechanically active proteins. Additionally, the current model was trained on unperturbed cells and becomes less accurate when asked to predict cellular phenotypes outside of the present training conditions. We suggest caution when using TensionDL to predict perturbations in cellular phenotypes associated with drug treatment or mutations. Training TensionDL to explicitly recognize the mechanical outputs of mutated or pharmacologically perturbed cells could be an excellent direction for future research, eventually enabling its use in drug or CRISPRi screens^34^. Additionally, our model currently excels at spatial predictions but is somewhat less accurate at quantitative predictions. TensionDL performs poorly (55% accuracy) at predicting which of two cells generates more tension fluorescence; however, it is spatially accurate, producing an average pixel-wise pearson of 0.83 across our entire test dataset. In contrast, the pixel-wise pearson of TensionDL-Q is lower, r=0.79, but it achieves 71% accuracy in a pairwise MTP fluorescence comparison test. This suggests that, at the current time, there is tradeoff between spatial accuracy and quantitative prediction. A larger training set which includes perturbations of cellular mechanics and future innovations in machine learning may overcome some of these difficulties.

We demonstrate TensionDL’s ability to measure forces on a heterogeneous surface on which both MTPs and ECM components are present. Future work should seek to extend TensionDL to native contexts. Force measurements in complex environments has previously been achieved via injection of oil bubbles^35^ or by genetically encoding tension sensors in an organism^36^. These or similar techniques may offer a route to validate TensionDL in native context. TensionDL’s compatibility with immunofluorescence detection of mechanically active proteins may offer a route to inferring force distributions in patient biopsies or histology workflows once the technology is sufficiently developed.

TensionDL aligns with the emerging field of *in silico* biophysics^34^. Deep learning has been employed to predict TFM maps^17^; however, our technique offers improved spatial resolution and pN sensitivity. Additionally, we demonstrate cell to cell mechanical comparisons which was not previously possible. Ultimately, integrating TensionDL with TFM may offer a route to augmenting both technologies, improving the spatial resolution of TFM by constraining maps with TensionDL or improving the quantitative interpretation of TensionDL via TFM measurements. Other approaches have sought to infer tissue stiffness from fluorescence images^37^. TensionDL offers the complementary capability of *in silico* receptor force measurement. In summary, we believe that TensionDL offers a method capable of assessing pN forces generated by cells from widely accessible imaging modalities. We anticipate that this technology will broaden access to MTP measurements which previously have been limited due to their complexity of use. Finally, the compatibility of TensionDL with standard fluorescence measurements promises future integration with high-throughput and clinical workflows.

## Materials and Methods

### Reagents

DNA oligonucleotides were custom-synthesized and HPLC-purified by Integrated DNA Technologies (IDT). Specific sequences are listed in **Supplementary Table 1**. cRGD-azide (Fisher Scientific, 50-168-6931) was used for CuAAC conjugation. EZ-Link NHS-Biotin (Thermo Fisher, PI20217) and Amine-PEG₂-Biotin (Thermo Fisher, 21346) were used for surface biotinylation. Cy3B-NHS ester was obtained from BroadPharm (BP28886). DMEM, fetal bovine serum (FBS), bovine calf serum (BCS), phosphate-buffered saline (PBS), TrypLE, penicillin–streptomycin, acetonitrile, ethanol, DMSO, triethylamine, trifluoroacetic acid, and Nanopure water were obtained from Thermo Fisher Scientific. Copper-click chemistry utilized CuSO₄, THPTA ligand, and sodium ascorbate (Thermo Fisher Scientific). Surface-modification reagents included APTES (Sigma Aldrich, 741442), 3-(trimethoxysilyl)propyl methacrylate (Sigma Aldrich, 216555000), and glacial acetic acid (Sigma Aldrich, 695092). 25-mm #1.5 glass coverslips (Thomas Scientific, 1217N82) and glass-bottom dishes (Cellvis, NC0409658) were used as substrates. Streptavidin was purchased from Rockland Immunochemicals (50-105-9142), and bovine serum albumin (BSA) from Fisher Bioreagents. Fibronectin (Sigma Aldrich, F1141) was used for ECM coating. HPLC purification was performed using an Agilent AdvanceBio Oligonucleotide C18 column, 4.6 × 150 mm (Agilent, 653950-702). Desalting steps used Amicon Ultra centrifugal filters, 3 kDa MWCO (Millipore, UFC500324). Hydrogel components included 40% acrylamide (Bio-Rad, 1610140), 2% bis-acrylamide (Bio-Rad, 1610142), acrylic acid (Santa Cruz, sc-358655), ammonium persulfate (APS; Bio-Rad, 1610700), and TEMED (Bio-Rad, 1610800). Activation reagents included EDC-HCl (Santa Cruz, sc-219152B), sulfo-NHS (Thermo Fisher, PI24510), and MES buffer components. Drug-treatment reagents included CK-666 (Arp2/3 inhibitor, Sigma Aldrich, 182515), Latrunculin B (Sigma Aldrich, L5288), and Y-27632 (Sigma Aldrich, product code). The mouse monoclonal anti-vinculin antibody (clone 7F9) was purchased from Thermo Fisher Scientific (53-9777-82). Additional consumables included 0.2-µm filtration units, Attofluor imaging chambers (Thermo Fisher, A7816), and other standard laboratory-grade solutions. All buffers were prepared using 18.2 MΩ Nanopure water.

### Tension probe synthesis

Probes were synthesized using a modification of previously reported methods^7,38^. The ligand strand (see **Supplementary Table 1**) carries a 5-octadiynyl-dU at the 5′ end and a 3′ primary amine, enabling sequential conjugation of cRGD-N_3_ and Cy3B-NHS (Supplementary Fig 1). The alkyne-modified DNA was first reacted with cRGD-azide via copper(I)-catalyzed azide-alkyne cycloaddition (CuSO₄ 100 µM, THPTA 500 µM, sodium ascorbate 1 mM, overnight at room temperature). The crude product was desalted using a 3 kDa Amicon centrifugal filter (10 min, 15,000 × g, 4 °C) and purified by reverse-phase HPLC using an Advanced Oligonucleotide C18 column using mobile-phase of 0.1 M triethylammonium acetate (TEAA) at pH 8 to acetonitrile gradient with 500 µL/min flow. The gradient was initiated at 10 % acetonitrile and increased by 0.5 % per minute; the cRGD–LigandDNA conjugate eluted at ∼21 min (**Supplementary Fig. 1a**). MALDI-TOF MS (linear mode, sinapinic acid matrix) confirmed successful coupling (Supplementary Fig. 1b). The purified cRGD–LigandDNA–NH₂ was subsequently labeled with Cy3B-NHS ester (0.1 M NaHCO₃, overnight at room temperature, protected from light), desalted (3 kDa filters), and purified by HPLC under identical conditions as listed above. The final cRGD–LigandDNA–Cy3B product eluted at ∼26.5 min and exhibited characteristic absorbance at 260 nm (DNA), and 560 nm (Cy3B) (**Supplementary Fig. 1c**). Probe concentrations were determined on a Nanodrop 2000 spectrophotometer using the theoretical extinction coefficient of the ligand strand, and aliquots were stored at 4 °C until use.

### Hairpin hybridization

Ligand, anchor, and hairpin-backbone oligonucleotides (sequences in **Supplementary Table 1**) were mixed at equimolar stoichiometry in 1× PBS to a final concentration of 300 nM in 0.2 mL low-bind Thermowell tubes. Mixtures were denatured at 90 °C for 5 min on a thermal cycler and annealed by cooling at 2 °C min⁻¹ to room temperature. Immediately after annealing, assemblies were used directly for surface immobilization by diluting 1:10 in PBS on the functionalized surface, yielding a 30 nM final strand concentration during immobilization.

### Cell culture

Mouse embryonic fibroblasts expressing GFP-vinculin^39^ were maintained in high-glucose DMEM supplemented with 10% BCS and 1% penicillin–streptomycin at 37 °C in a humidified 5% CO₂ incubator. NIH/3T3 mouse fibroblasts (CRL-1658, ATCC) and MDCK II canine kidney epithelial cells (CRL-2936, ATCC) were cultured in high-glucose DMEM supplemented with 10% FBS and 1% penicillin–streptomycin under identical conditions. Cells were passaged at ∼80% confluency using standard trypsinization. For experiments, cells were rinsed once with PBS, detached with trypsin for 3 min, pelleted at 400 × g for 5 min, and resuspended at 1.4 × 10⁶ cells mL⁻¹. Approximately 7 × 10⁴ cells were seeded onto each prepared functionalized surface for downstream experiments.

### Surface preparation

Surfaces were prepared with minor modification to previously described protocols^7^. Briefly, coverslips were sonicated in Nanopure water (3 × 3 min), then in ethanol (3 × 3 min), and dried at 100 °C for 15 min. Substrates were cleaned in piranha solution (3: 1 mixture of H₂SO₄ to 30% H₂O₂) for 15 min (Caution: piranha solution is extremely corrosive and may explode if mixed with organic substances). After piranha cleaning, surfaces were rinsed thoroughly with Nanopure water (5 washes), sonicated for 5 min in water, and base-etched in 1 M NaOH for 1 h. After five water washes and three ethanol washes, surfaces were silanized in 3% (v/v) APTES in ethanol for 1 h, rinsed in ethanol (3 washes), dried under a stream of N₂, and cured at 100 °C for 1 h. Once cooled, amine-terminated surfaces were biotinylated overnight with Biotin-NHS (5 mg/mL in DMSO). The next day, coverslips were rinsed in ethanol (3 washes), dried under a stream of N₂, mounted into Attofluor imaging chambers (ThermoFisher, A7816), washed with PBS (3 × 5 mL), blocked with 5% BSA in PBS for 30 min, and washed again with PBS (3 × 5 mL). Streptavidin (10 µg/mL in PBS) was applied for 1 h at room temperature, followed by three PBS washes. Hybridized DNA tension probes were then introduced at 30 nM in PBS (prepared as 300 nM assemblies diluted 1:10 directly on the surface) and incubated for 1 h at room temperature, protected from light, prior to imaging.

### Image acquisition

Fluorescence imaging was performed on a Nikon Eclipse Ti2 inverted microscope equipped with ring-TIRF illumination (ILAS Ring-TIRF) and a 100×/1.49 NA oil-immersion Apo objective (Nikon Instruments). Images were recorded on a QUEST CMOS camera using NIS-Elements. To minimize channel bleed-through, we used Nikon filter modules: for the 561-nm channel, a 595/31 nm emission filter; for the 488-nm channel, a 515/30 emission filter. Excitation, exposure, and acquisition settings are provided in **Supplementary Table 2**. Within each experiment, illumination and detection settings were held constant and channels were acquired sequentially.

### Training dataset Image processing

For each field of view, three images were acquired: MTP tension fluorescence, RICM, and GFP–vinculin fluorescence. Raw tension images underwent preprocessing: first, a constant camera offset of 215 intensity units was subtracted (dark-signal correction), then flat-field correction was applied by normalizing each pixel by the median-derived illumination profile to remove the uneven illumination profile of the laser. 256 × 256 pixel square ROIs were manually selected. For each accepted ROI, cropped RICM (input), vinculin (input), and tension (target) were saved.

### TensionDL training

A total of 1312 ROIs (256 × 256 pixels) were manually extracted from cells across 6 independent experiments and randomly partitioned into training, validation, and test sets. To improve model robustness, 30 and 34 cell-free background ROIs were added to the training and validation datasets, respectively. The model was implemented using CytoDL v0.1.8 (github.com/AllenCell/cyto-dl) with a U-Net encoder-decoder architecture featuring 18 convolutional layers (3×3 kernels), four downsampling layers (stride-2 convolutions), four upsampling layers (2×2 transposed convolutions), and skip connections between corresponding encoder and decoder levels (**Supplementary Fig. 5**). Training was performed on an NVIDIA A100 GPU using 128 × 128 pixel patches extracted from the input ROIs, with a batch size of 64. The model was optimized using the Adam optimizer (learning rate: 0.01, weight decay: 1×10⁻⁴, β=[0.5, 0.999]) with exponential learning rate decay (γ=0.995), ReLU activation functions, InstanceNorm normalization, and mean squared error loss over 1000 epochs. Inference utilized a sliding window approach with 25% overlap and Gaussian blending. The trained model and training data will be publicly available upon publication.

### Image segmentation and Subcellular Analysis

TensionDL and GT images were segmented in MATLAB 2024a. Briefly, images were normalized, subjected to an adaptive intensity threshold using the function “imbinarize,” and then subjected to Chan-vese active contouring. 4-connected pixel regions smaller than 60 pixels were removed to generate the final mask. 8-connected pixel regions were then identified with the function bwlabel. A representative example of the results of this segmentation can be seen in **Fig. 2a**. The jaccard similarity matrix was then computed for all identified tension zones (**Fig. 2b**). In cases where predicted tension zone or GT tension zone corresponded to multiple segmented regions, these zone were relabeled with the same index and the jaccard similarity matrix was recomputed for the purpose of the analyses in **Fig. 2**.

### Radial and angular intensity analysis

MTP tension maps were background-corrected by subtracting the average intensity in a manually drawn cell-free ROI. The standard deviation of the background signal was computed, and MTP fluorescence was then thresholded at two standard deviations above the background signal to create binary masks. Centroids were computed for each masked cell in both ground truth and predictions. For the radial analysis, the integrated MTP or TensionDL intensity was computed within each 30 pixel rings from the centroid to the farthest nonzero pixel. Ring intensities were divided by ring area to account for increasing ring area as a function of radius. Radial intensities were normalized to the max integrated ring intensity to facilitate cell to cell profile comparison. For the angular analysis, the integrated MTP and TensionDL intensities were computed in twelve equal sectors radiating from the centroid (30° each). To facilitate cell to cell comparison, angular intensities were normalized to the max integrated angular intensity observed for each cell. Agreement between ground truth and predictions was assessed using Kuiper’s two-sample t-test on the cumulative distributions of the radial and angular profiles; p < 0.05 indicated significantly different spatial patterns.

### Hydrogel preparation

Poly(acrylamide-co-acrylic acid) hydrogels of defined stiffness were prepared by varying the acrylamide to bis-acrylamide ratio while maintaining a constant acrylic acid concentration to permit subsequent bioconjugation. Glass-bottom dishes were functionalized with bind-silane by incubating them in a solution of glacial acetic acid and 3-(trimethoxysilyl)propyl methacrylate in deionized water for 30 min, followed by two 15-min rinses in Milli-Q water. Precursor solutions were prepared by combining acrylamide and bis-acrylamide at ratios listed in **Supplementary Table 4**, adding acrylic acid and 10× PBS to a final volume of 500 µL, and adjusting the pH to 7.5–8.0 with NaOH. After degassing for 30 min, polymerization was initiated with ammonium persulfate (APS) and TEMED. A 10 µL aliquot of precursor was dispensed onto each silanized dish and overlaid with a Rain-X–treated coverslip to form thin, uniform gels. Following polymerization, coverslips were released by adding PBS, and hydrogels were stored overnight at 4 °C in PBS before use.

### Hydrogel mechanical characterization

Hydrogel stiffness was measured using a stress-controlled rheometer equipped with a 25 mm parallel-plate geometry. The storage modulus (G′) was recorded at 1% strain and 1 Hz and averaged over 50 readings. Young’s modulus (E) was calculated from G′ and averaged across at least three gels per formulation.

### Conjugation of MTP to hydrogels

Hydrogels were rinsed with PBS before activation of surface carboxyl groups in EDC/sulfo-NHS [conc] activation buffer for 20 min. Activated gels were washed with PBS and incubated for 2 h with Amine-PEG₂-Biotin diluted in PBS. After biotinylation, gels were rinsed, blocked with 5% BSA for 30 min, and washed again. Molecular tension probe assemblies (60 nM) were mixed with streptavidin (10ug/ml) then applied to the surface for 1 h before subsequent experiments.

### Fixation and immunofluorescence staining

To preserve the MTP signal during fixation and subsequent immunostaining, a DNA locking protocol was implemented^23^. The fixation and permeabilization process can disrupt MTP signal; therefore, locking was employed to stabilize the opened hairpin conformation prior to fixation. Cells were allowed to adhere and spread on MTP-functionalized surfaces for 15 minutes at 37°C with 5% CO₂. The locking strand (100nM in PBS) was then added directly to cells and incubated for 1 minute at room temperature. This brief incubation time was sufficient for strand hybridization to the force-opened hairpins while minimizing the effects of cell dynamics. Following the 1-minute incubation, cells were washed once with PBS and immediately fixed. Cells were fixed in 4% paraformaldehyde (PFA) in phosphate-buffered saline (PBS) for 15 minutes at room temperature. Following three washes with PBS, cells were permeabilized with 0.1% Triton X-100 in PBS for 10 minutes and blocked with 1% bovine serum albumin (BSA) in PBS for 1 hour at room temperature. Samples were incubated with mouse anti-vinculin antibody (1:100 dilution in PBS) for 1 hour at RT. After washing three times with PBS, cells were imaged.

### Fibronectin MTP surface preparation

Following MTP coating, surfaces were rinsed once with PBS. Fibronectin was added to the MTP-coated surfaces at a final surface concentration of 0.5-5 μg/ml in PBS. The fibronectin solution was incubated for 60 minutes at 37°C. After fibronectin incubation, surfaces were washed once with PBS and used immediately for cell seeding and imaging.

### Fibronectin timelapse analysis

Cell footprints were segmented from RICM images, centroids were computed, and boundaries were tracked in each frame; early and late outlines were overlaid to illustrate edge motion. For angular summaries, images were divided into evenly spaced angle bins around the centroid. Angle-wise predicted MTP intensity (background-subtracted and frame-normalized) was plotted at single time points as angular histograms. To visualize changes over time, angular histograms were also displayed as a kymograph. A similar analysis was performed to assess cell surface contact area from the RICM-derived masks.

### Cytoskeletal inhibitor treatments

Drug perturbations were applied during live cell timelapse imaging. Cells were allowed to spread on surfaces for 15 minutes and imaged at 3 minute time intervals for 40 minutes. At 5 minutes after beginning timelapse imaging, compounds were added directly to the imaging medium at the following final concentrations: CK-666 (150 µM) and Latrunculin B (1 µM), each prepared from DMSO stock solutions, and Y-27632 (5 µM) prepared from an aqueous stock. For pairwise comparison between before and after drug treatment, each cell was normalized by the max integrated intensity for that cell during the timelapse in order to account for cell to cell and experimental variability. A matched vehicle-only control (DMSO) was imaged and analyzed under identical conditions (**Supplementary Fig. 13**).

### UMAP Analysis

Raw images were normalized to [0,1] intensity range and each grayscale image was then converted to RGB format (by replicating across three channels) to be compatible with the pre-trained DINOv2 vision transformer^40^, which extracted 384-dimensional feature vectors from each image. These feature vectors captured complex morphological and textural patterns. The high-dimensional features were then compressed into 2D coordinates using UMAP dimensionality reduction, allowing visualization of the entire dataset. Final visualizations displayed these UMAP embeddings color-coded by experimental condition (fibronectin concentration) to reveal treatment-specific clustering patterns (**Supplementary Fig 9**).

### TensionDL-Q Model

All images were first background-normalized by drawing a uniform, cell-free ROI per frame and dividing the entire image by the mean intensity of that ROI, yielding a common scale (∼1–25 a.u.). After this step, we computed the dataset-wide median and maximum pixel intensities across the training set. Intensity standardization was then applied using an intensity-normalization transform that (i) subtracts the dataset median from every pixel and (ii) divides by the dataset maximum, producing inputs in the 0–1 range for the MTP channel. Outputs were directly saved as 32-bit floating-point values without normalization to preserve continuous intensity information.

## Acknowledgements

MEF GFP-vinculin cells were a gift from A. Garcia. J. Brockman would like to acknowledge support from R35GM157007. We acknowledge D. Mehrotra for helpful discussion. D. Smith would like to acknowledge support from T32GM158463.

## References

1. Zhang, Y. et al. Platelet integrins exhibit anisotropic mechanosensing and harness piconewton forces to mediate platelet aggregation. Proc. Natl. Acad. Sci. U. S. A. 115, 325–330 (2018).

2. Engler, A. J., Sen, S., Sweeney, H. L. & Discher, D. E. Matrix elasticity directs stem cell lineage specification. Cell 126, 677–689 (2006).

3. Liu, Y. et al. DNA-based nanoparticle tension sensors reveal that T-cell receptors transmit defined pN forces to their antigens for enhanced fidelity. Proc. Natl. Acad. Sci. U. S. A. 113, 5610–5615 (2016).

4. Sibener, L. V. et al. Isolation of a Structural Mechanism for Uncoupling T Cell Receptor Signaling from Peptide-MHC Binding. Cell 174, 672–687.e27 (2018).

5. Grashoff, C. et al. Measuring mechanical tension across vinculin reveals regulation of focal adhesion dynamics. Nature 466, 263–266 (2010).

6. Stabley, D. R., Jurchenko, C., Marshall, S. S. & Salaita, K. S. Visualizing mechanical tension across membrane receptors with a fluorescent sensor. Nat. Methods 9, 64–67 (2011).

7. Zhang, Y., Ge, C., Zhu, C. & Salaita, K. DNA-based digital tension probes reveal integrin forces during early cell adhesion. Nat. Commun. 5, 5167 (2014).

8. Brockman, J. M. et al. Live-cell super-resolved PAINT imaging of piconewton cellular traction forces. Nat. Methods 17, 1018–1024 (2020).

9. Liu, Y., Galior, K., Ma, V. P.-Y. & Salaita, K. Molecular tension probes for imaging forces at the cell surface. Acc. Chem. Res. 50, 2915–2924 (2017).

10. Saraswathibhatla, A., Indana, D. & Chaudhuri, O. Cell-extracellular matrix mechanotransduction in 3D. Nat. Rev. Mol. Cell Biol. 24, 495–516 (2023).

11. Panciera, T. et al. Reprogramming normal cells into tumour precursors requires ECM stiffness and oncogene-mediated changes of cell mechanical properties. Nat. Mater. 19, 797–806 (2020).

12. Isola, P., Zhu, J.-Y., Zhou, T. & Efros, A. A. Image-to-image translation with conditional adversarial networks. in 2017 IEEE Conference on Computer Vision and Pattern Recognition (CVPR) 5967–5976 (IEEE, 2017).

13. Rivenson, Y. et al. Virtual histological staining of unlabelled tissue-autofluorescence images via deep learning. Nat Biomed Eng 3, 466–477 (2019).

14. Bai, B. et al. Deep learning-enabled virtual histological staining of biological samples. Light Sci Appl 12, 57 (2023).

15. Ounkomol, C., Seshamani, S., Maleckar, M. M., Collman, F. & Johnson, G. R. Label-free prediction of three-dimensional fluorescence images from transmitted-light microscopy. Nat. Methods 15, 917–920 (2018).

16. Christiansen, E. M. et al. In silico labeling: Predicting fluorescent labels in unlabeled images. Cell 173, 792–803.e19 (2018).

17. Schmitt, M. S. et al. Machine learning interpretable models of cell mechanics from protein images. Cell 187, 481–494.e24 (01/2024).

18. Sabass, B., Gardel, M. L., Waterman, C. M. & Schwarz, U. S. High resolution traction force microscopy based on experimental and computational advances. Biophys. J. 94, 207–220 (2008).

19. Jaccard, P. Nouvelles recherches sur la distribution florale. Bulletin de la Societe Vaudoise des Sciences Naturelles 44, 223–270 (1908).

20. Caicedo, J. C. et al. Nucleus segmentation across imaging experiments: the 2018 Data Science Bowl. Nat. Methods 16, 1247–1253 (2019).

21. Discher, D. E., Janmey, P. & Wang, Y.-L. Tissue cells feel and respond to the stiffness of their substrate. Science 310, 1139–1143 (2005).

22. Wang, W. et al. Hydrogel-based molecular tension fluorescence microscopy for investigating receptor-mediated rigidity sensing. Nat. Methods 20, 1780–1789 (2023).

23. Ma, R. et al. DNA probes that store mechanical information reveal transient piconewton forces applied by T cells. Proc. Natl. Acad. Sci. U. S. A. 116, 16949–16954 (2019).

24. Vroemen, P. A. M. M. et al. The importance of coating surface and composition for attachment and survival of neuronal cells under mechanical stimulation. J. Biomed. Mater. Res. A 113, e37919 (2025).

25. Ronneberger, O., Fischer, P. & Brox, T. U-Net: Convolutional Networks for Biomedical Image Segmentation. arXiv [cs.CV] (2015) doi:10.48550/arXiv.1505.04597.

26. Jurchenko, C. & Salaita, K. S. Lighting up the force: Investigating mechanisms of mechanotransduction using fluorescent tension probes. Mol. Cell. Biol. 35, 2570–2582 (2015).

27. Ringer, P. et al. Multiplexing molecular tension sensors reveals piconewton force gradient across talin-1. Nat. Methods 14, 1090–1096 (2017).

28. LaCroix, A. S., Rothenberg, K. E., Berginski, M. E., Urs, A. N. & Hoffman, B. D. Construction, imaging, and analysis of FRET-based tension sensors in living cells. Methods Cell Biol. 125, 161–186 (2015).

29. Borghi, N. et al. E-cadherin is under constitutive actomyosin-generated tension that is increased at cell-cell contacts upon externally applied stretch. Proc. Natl. Acad. Sci. U. S. A. 109, 12568–12573 (2012).

30. Butler, J. P., Tolić-Nørrelykke, I. M., Fabry, B. & Fredberg, J. J. Traction fields, moments, and strain energy that cells exert on their surroundings. Am. J. Physiol. Cell Physiol. 282, C595–605 (2002).

31. del Álamo, J. C. et al. Three-dimensional quantification of cellular traction forces and mechanosensing of thin substrata by fourier traction force microscopy. PLoS One 8, e69850 (2013).

32. Plotnikov, S. V., Sabass, B., Schwarz, U. S. & Waterman, C. M. High-resolution traction force microscopy. Methods Cell Biol. 123, 367–394 (2014).

33. Schwarz, U. S. & Soiné, J. R. D. Traction force microscopy on soft elastic substrates: A guide to recent computational advances. Biochim. Biophys. Acta 1853, 3095–3104 (2015).

34. Oria, R., Jain, K. & Weaver, V. M. Exploring the intersection of mechanobiology and artificial intelligence. NPJ Biol. Phys. Mech. 2, 9 (2025).

35. Campàs, O. et al. Quantifying cell-generated mechanical forces within living embryonic tissues. Nat. Methods 11, 183–189 (2014).

36. Lemke, S. B., Weidemann, T., Cost, A.-L., Grashoff, C. & Schnorrer, F. A small proportion of Talin molecules transmit forces at developing muscle attachments in vivo. PLoS Biol. 17, e3000057 (2019).

37. Stashko, C. et al. A convolutional neural network STIFMap reveals associations between stromal stiffness and EMT in breast cancer. Nat. Commun. 14, 3561 (2023).

38. Brockman, J. M. et al. Mapping the 3D orientation of piconewton integrin traction forces. Nat. Methods 15, 115–118 (2018).

39. Zhou, D. W., Lee, T. T., Weng, S., Fu, J. & García, A. J. Effects of substrate stiffness and actomyosin contractility on coupling between force transmission and vinculin-paxillin recruitment at single focal adhesions. Mol. Biol. Cell 28, 1901–1911 (2017).

40. Oquab, M., et al. DINOv2: Learning robust visual features without supervision. arXiv [cs.CV] (2023).

