## Supplementary Information for "Prediction of Piconewton Receptor Tension Images using Deep Learning"

### Supplementary Figures

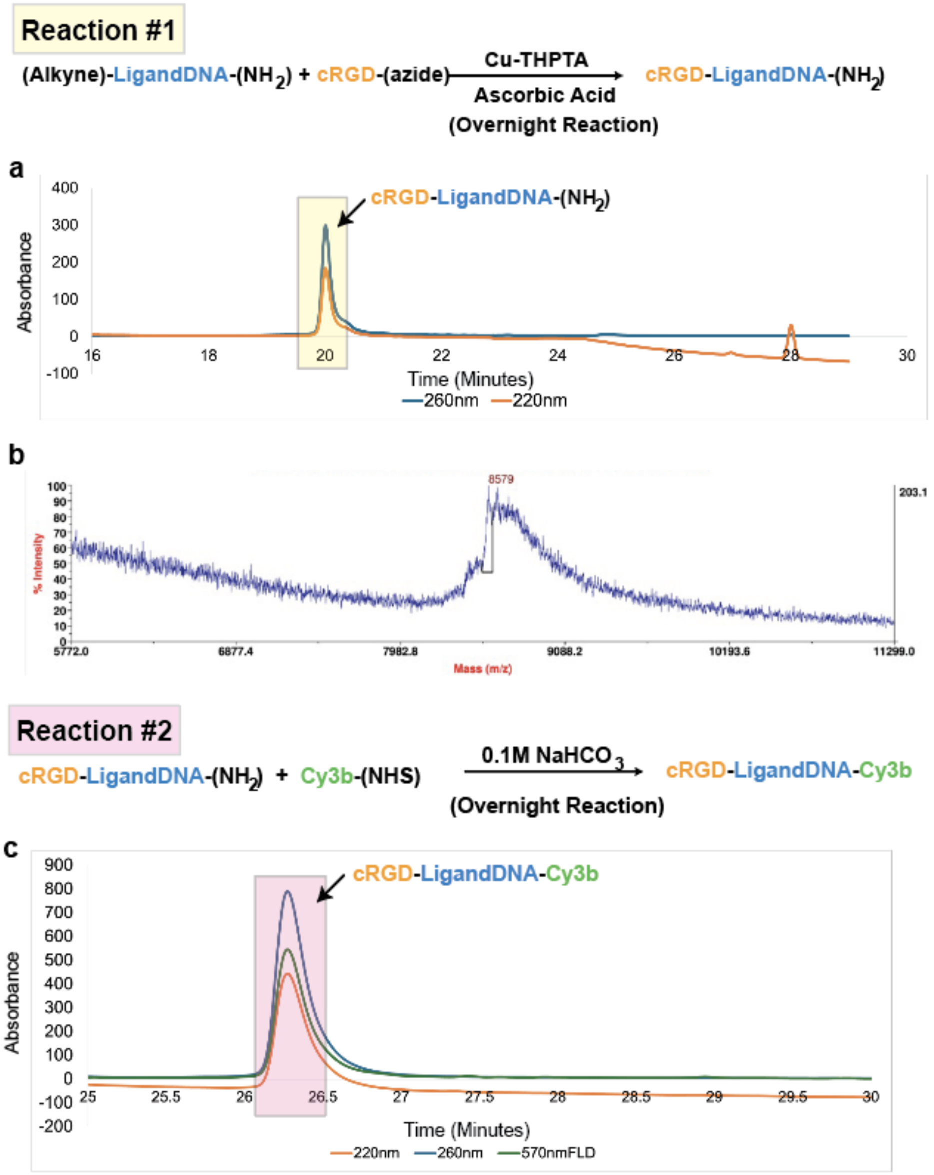


**Supplementary Figure 1: Synthesis and analytical characterization of cRGD–DNA MTP probes.** Reaction #1: Cu-THPTA/ascorbate-mediated azide–alkyne cycloaddition of (alkyne)–LigandDNA–($NH_2$) with cRGD–azide to give cRGD–LigandDNA–($NH_2$) (overnight). a, Analytical RP-HPLC chromatograms monitored at 220 and 260 nm, highlighting the cRGD–LigandDNA–($NH_2$) fraction. b, Mass spectrum of cRGD–LigandDNA–($NH_2$). Reaction #2: NHS-ester coupling of Cy3B–(NHS) to cRGD–LigandDNA–($NH_2$) in 0.1 M $NaHCO_3$ to create cRGD–LigandDNA–Cy3B (overnight). c, Analytical RP-HPLC chromatograms of the product monitored at 220 nm, 260 nm, and Cy3B fluorescence (~570 nm FLD).

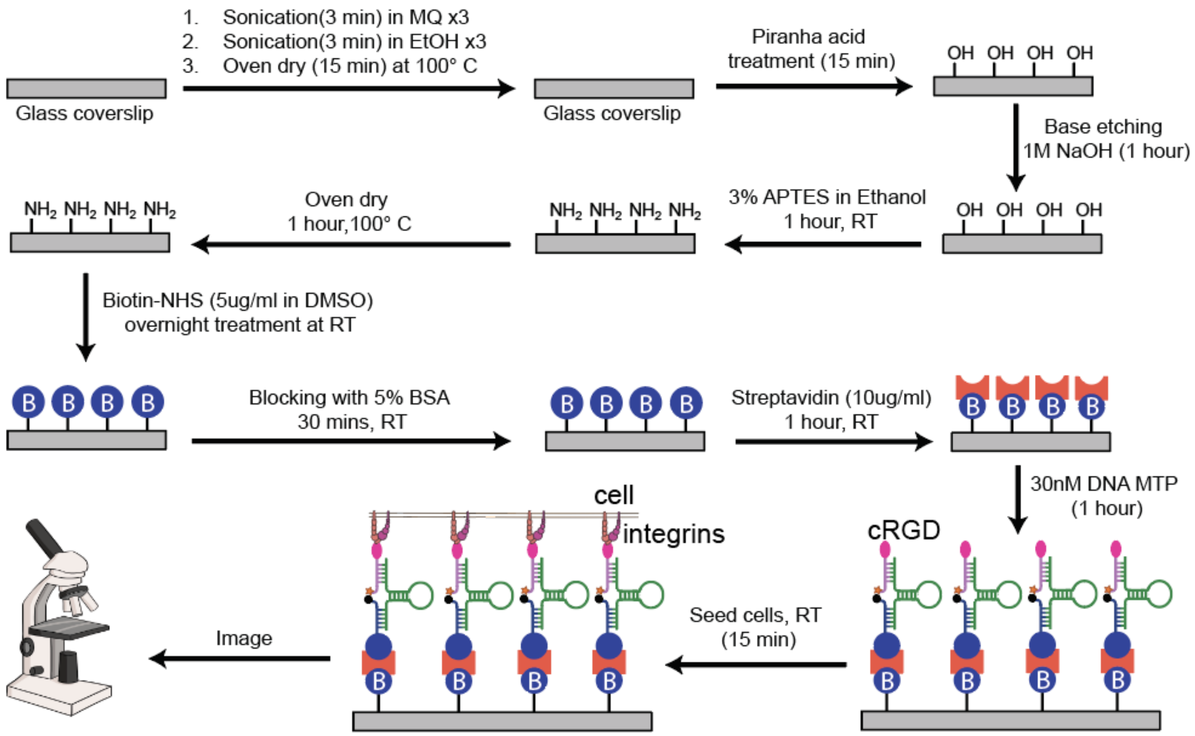


**Supplementary Figure 2: Workflow for functionalizing glass coverslips with cRGD–DNA MTPs**. Glass coverslips are cleaned (sonication in MQ ×3, EtOH ×3) and dried, then activated by piranha (15 min) and base etched in 1 M NaOH (1 h). Surfaces are silanized with 3% APTES in ethanol (1 h, RT) and cured at 100 °C. Primary amines are biotinylated with Biotin–NHS (5 µg/mL in DMSO, overnight, RT), blocked with 5% BSA (30 min, RT), and incubated with streptavidin (10 µg/mL, 1 h, RT). Finally, biotinylated DNA MTP (30 nM) presenting cRGD is immobilized (1 h), cells are seeded (15 min, 37° C) to allow integrin–cRGD engagement, and samples are imaged.

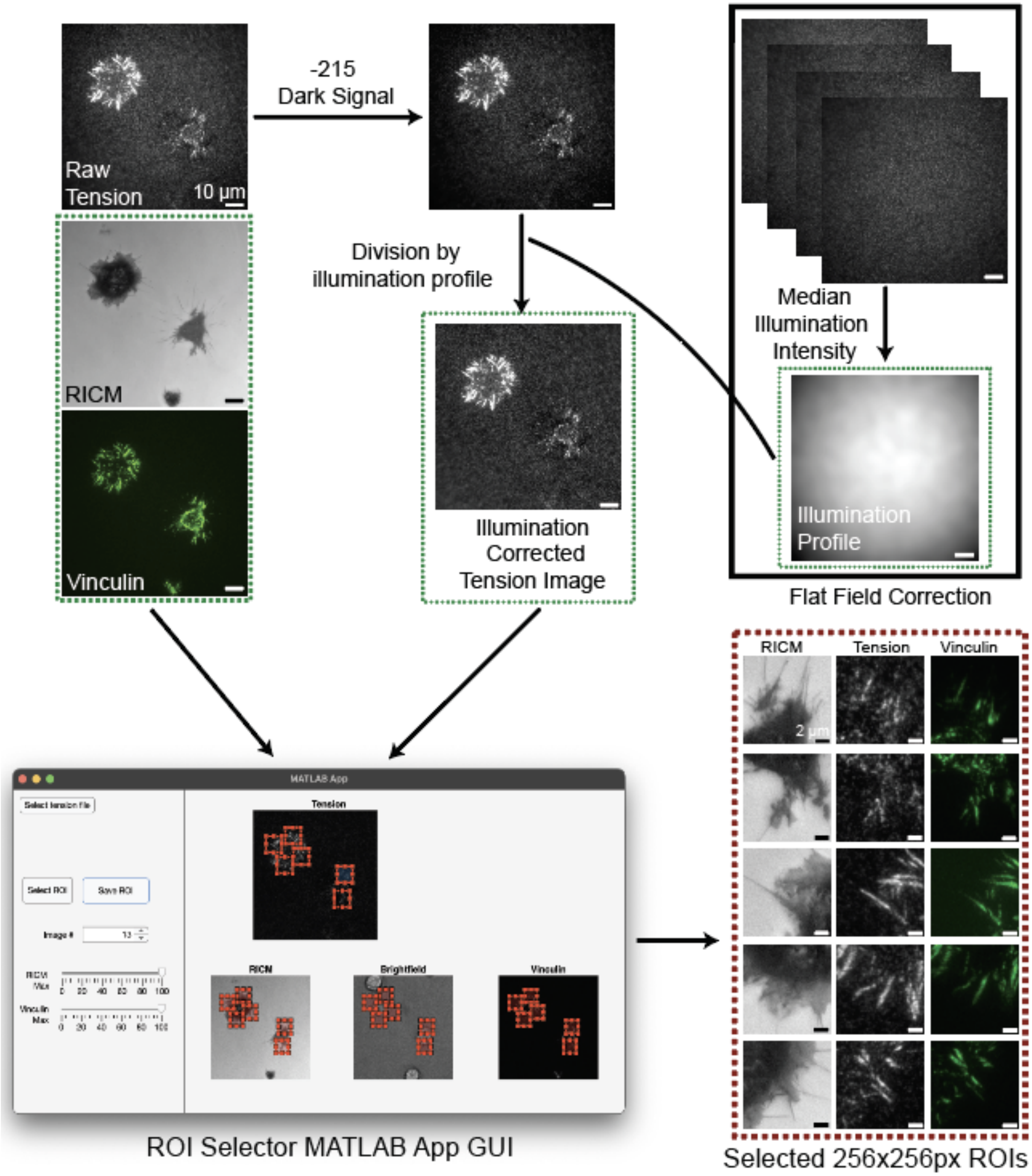


**Supplementary Figure 3: Dataset generation and preprocessing for TensionDL.** Raw MTP tension frames (scale bar, 10 µm) with paired RICM and GFP–vinculin images are first dark-corrected (subtracted 215 offset) and flat-fielded by dividing by a median illumination profile computed from image stacks. The illumination-corrected tension image, together with RICM and vinculin, is loaded into a custom ROI-Selector MATLAB app, where single-cell, non-overlapping 256 × 256 px ROIs are selected. Right: examples of the RICM / Tension / Vinculin crops used as model inputs/targets.

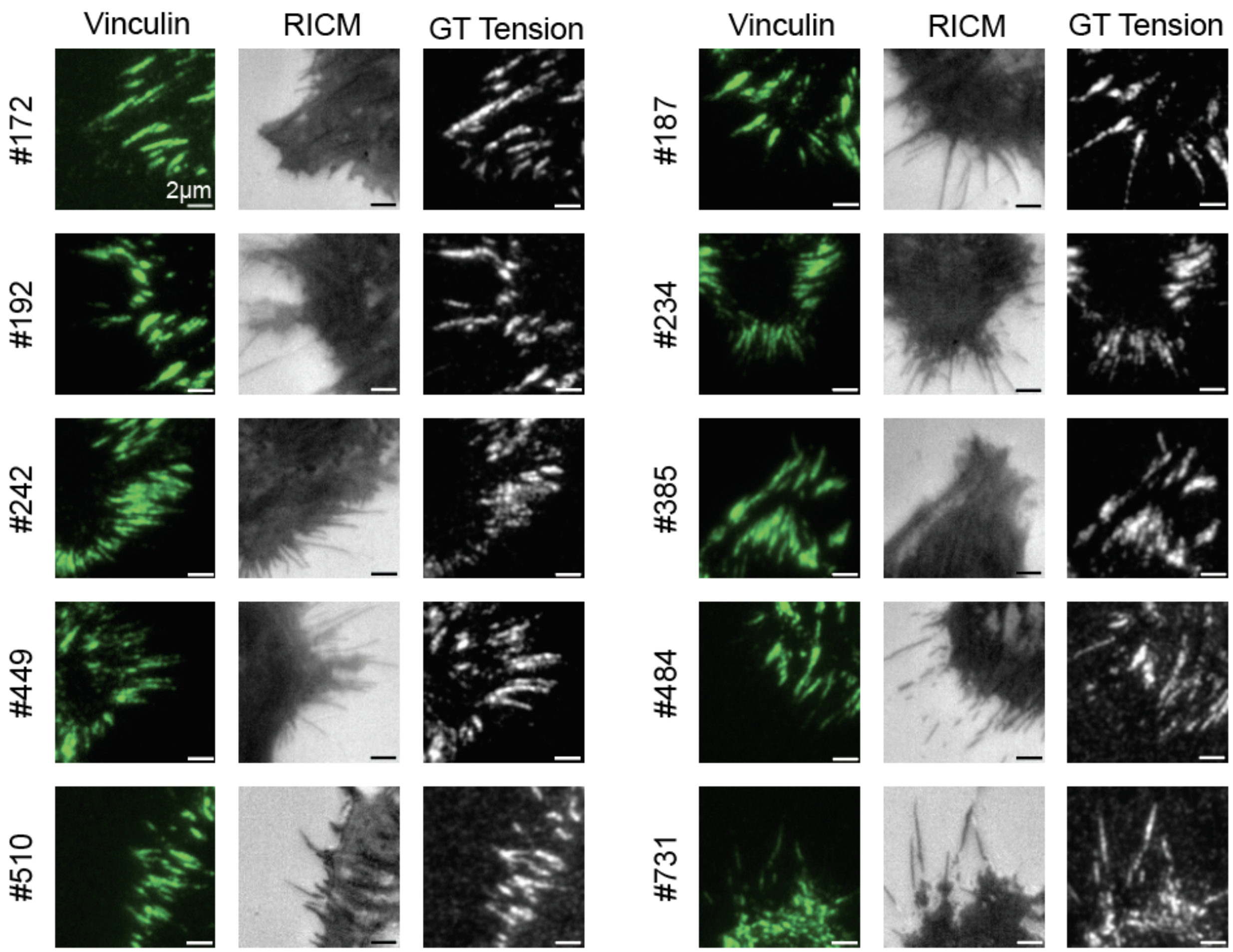


**Supplementary Figure 4: Representative training samples for TensionDL.**
Example 256 × 256 px ROIs from the training set, GFP–vinculin (left), RICM (middle), and ground-truth MTP-tension (right). Row tags indicate the original cell IDs. Scale bars, 2 μm.

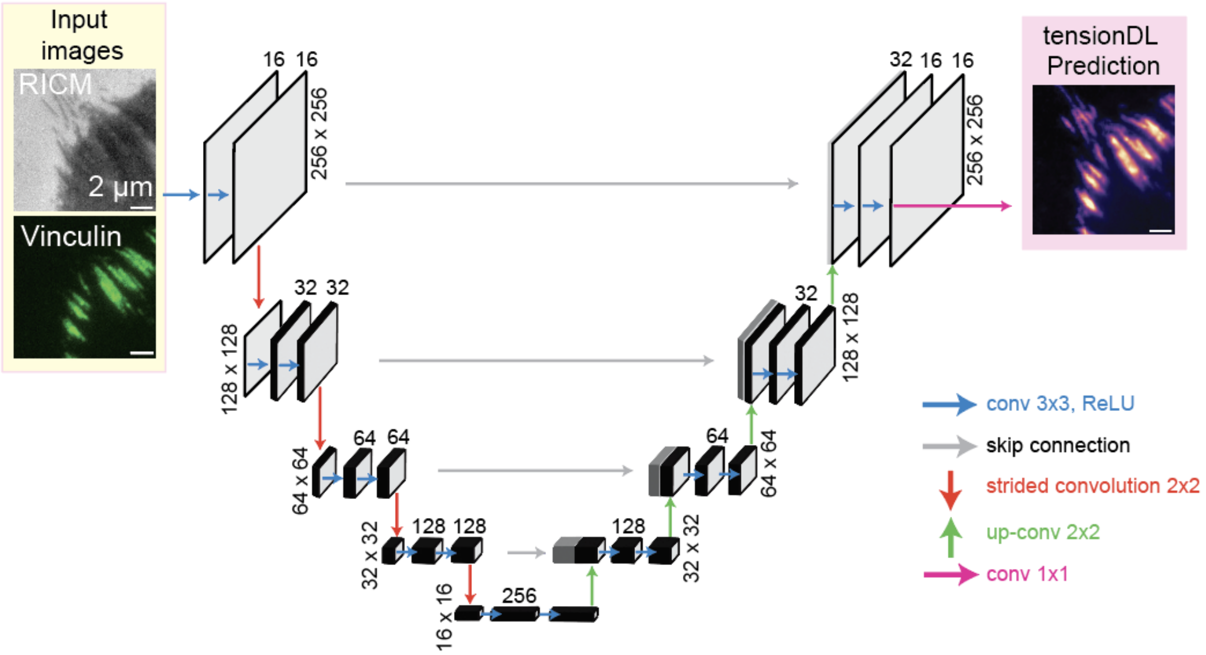


**Supplementary Figure 5: TensionDL model architecture.**
Schematic of the U-Net style encoder-decoder used to predict MTP-tension maps from paired RICM and GFP–vinculin inputs (256×256 px ROIs; scale bars, 2 µm). Convolutional blocks use 3×3 conv + ReLU (blue), with 2×2 strided convolutions for downsampling (red) and 2×2 transposed convolutions for upsampling (green). Skip connections concatenate encoder features to decoder features at matching resolutions. Numbers indicate feature-channel widths (16→32→64→128→256 and back). A final 1×1 convolution (magenta) produces a single-channel TensionDL prediction.

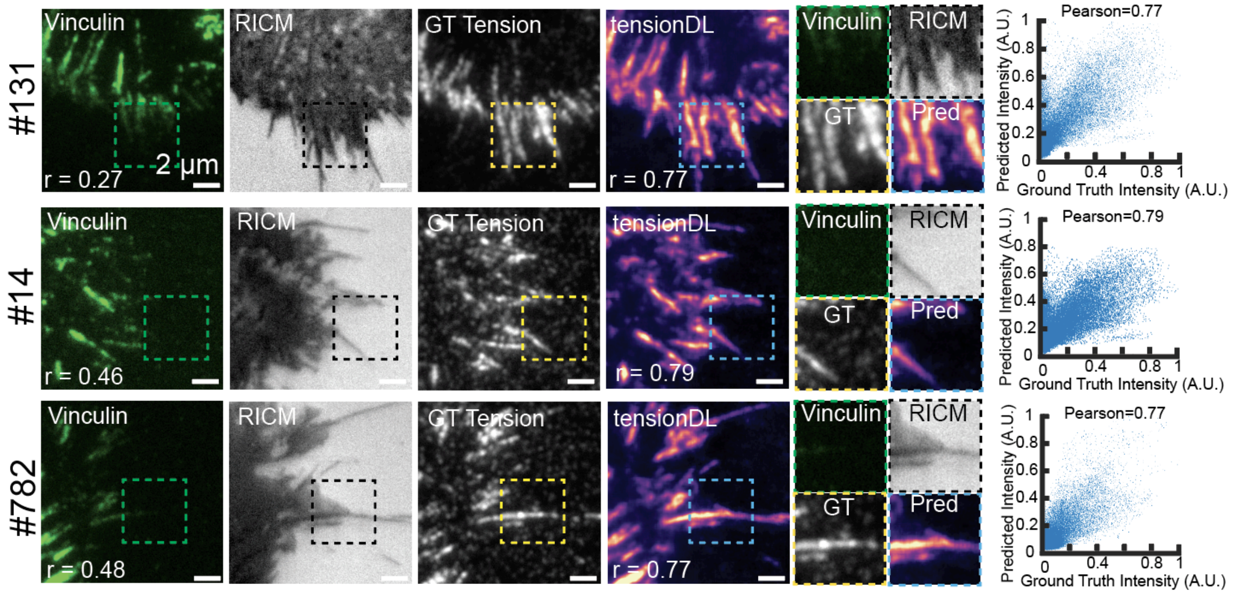


**Supplementary Figure 6: Additional examples of TensionDL predictions.**
For three test ROIs (cell IDs indicated at left), columns show GFP–vinculin, RICM, ground-truth (GT) MTP-tension, and TensionDL prediction. Dashed boxes mark regions enlarged to compare GT and Pred; several areas exhibit weak/absent vinculin contrast yet retain clear tension signals in both ground truth and prediction. Right: per-pixel scatter plots of Pred vs GT integrated intensity with Pearson *r* annotated. Note that the Pearson in Vinculin images is the pearson between Vinculin and Ground Truth while the pearson in the TensionDL image is the pearson between TensionDL prediction and GT. Scale bars, 2 µm.

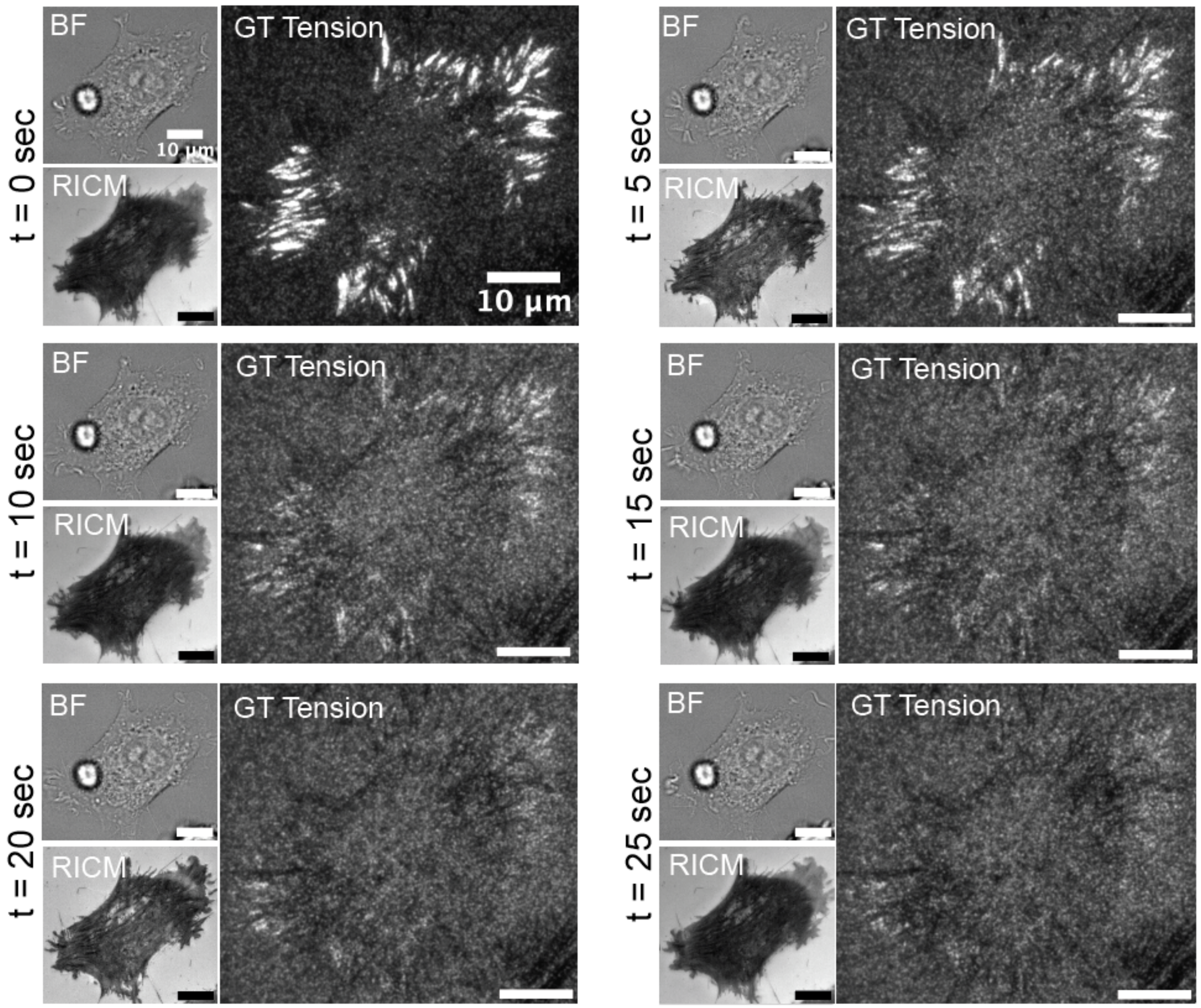


**Supplementary Figure 7: Photobleaching of DNA MTP tension probes during imaging**. Time-lapse GT MTP-tension images from a single cell acquired under continuous illumination at t = 0, 5, 10, 15, 20, and 25 s. Brightfield (BF) and RICM also shown for reference. The gradual loss of GT signal over time illustrates probe photobleaching. Scale bar, 10 µm.

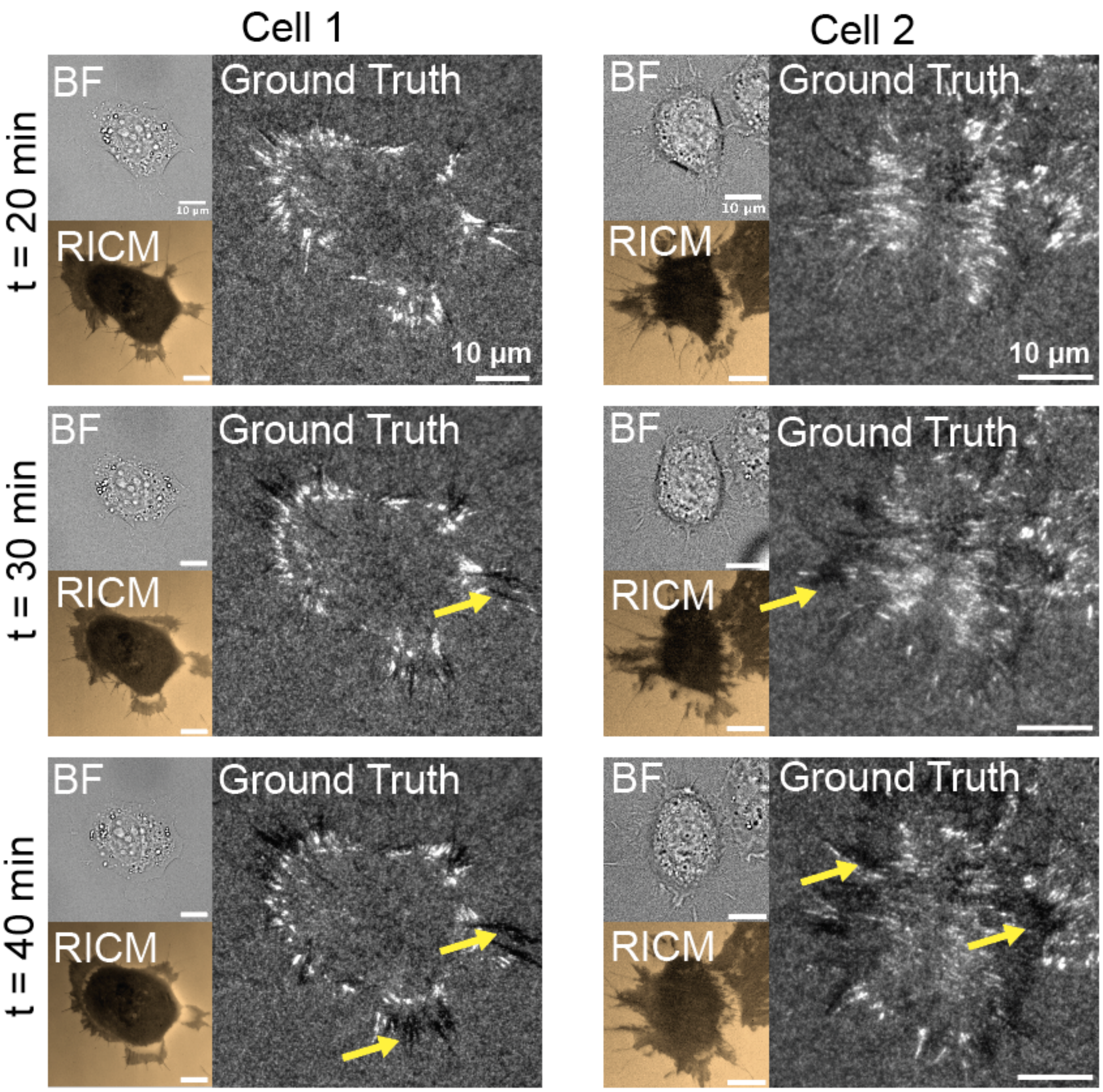


**Supplementary Figure 8: Cells rupture MTPs over a short time scale**
Representative time-lapse sequences from two individual cells showing progressive rupture of molecular tension probes (MTPs). For each time point (20, 30, and 40 min), brightfield (BF) and RICM images are shown alongside the corresponding ground-truth MTP signal. Yellow arrows highlight regions where localized black spots indicate probe rupture under sustained receptor tension. Scale bar, 10 μm.

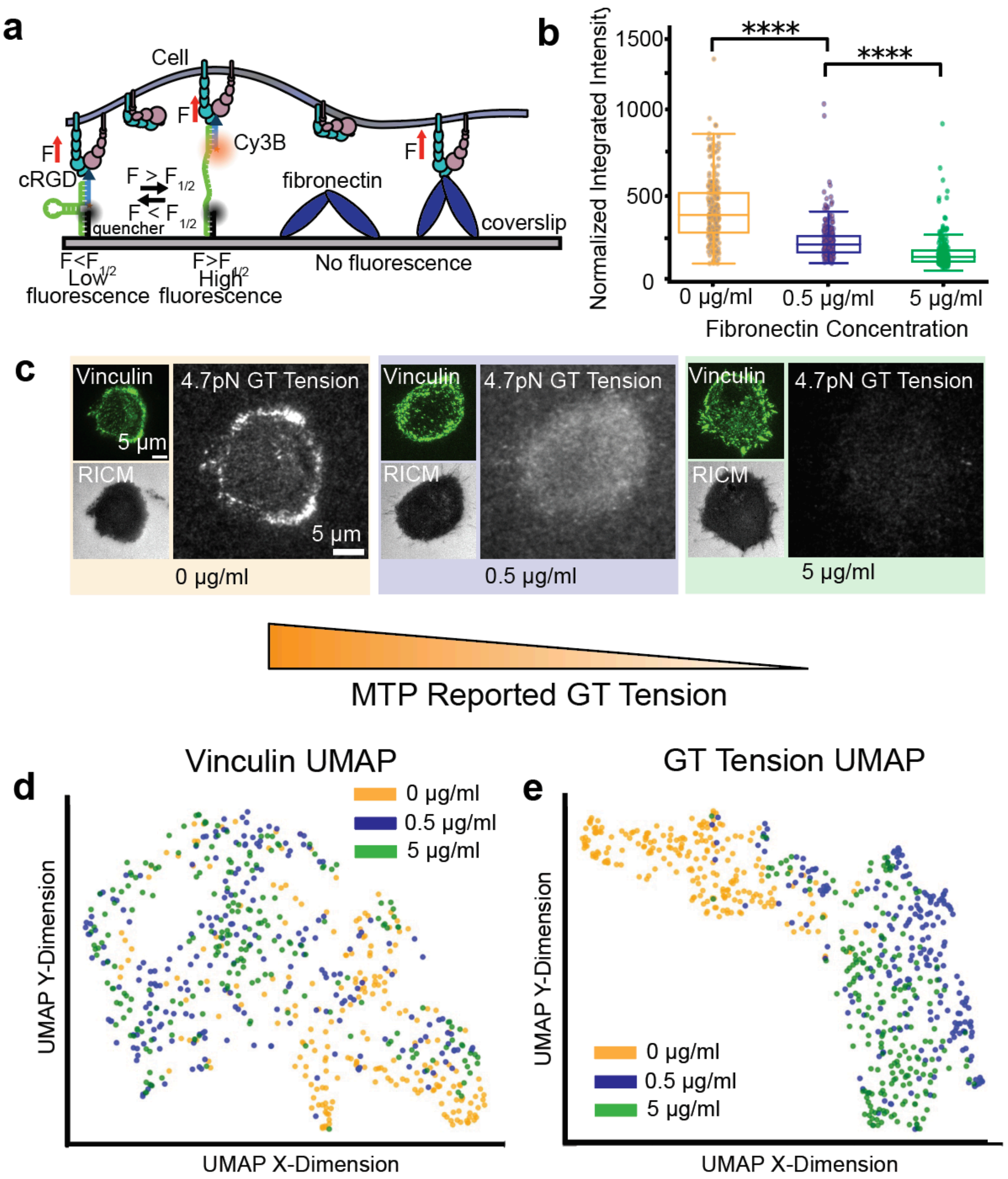


**Supplementary Figure 9: Effects of fibronectin concentration on MTP-reported forces**
**a.** Schematic of a coverslip coated with Fibronectin and co-functionalized with MTPs. Forces transmitted through fibronectin are not reported by the MTPs. **b.** Quantification of normalized integrated MTP intensity for cells seeded on 0 (231 cell ROIs), 0.5 (224 cell ROIs), and 5 (234 cell ROIs) µg/mL fibronectin surfaces co-functionalized with MTPs (**** $P < 0.0001$, Welch's t-test, n=3 independent experiments). **c.** Representative vinculin (left) and ground-truth 4.7 pN MTP tension images (right) from cells plated on surfaces coated with 0, 0.5, or 5 µg/mL fibronectin. **d.** UMAP embedding of vinculin features across conditions, color-coded by fibronectin concentration. **e.** UMAP embedding of ground-truth MTP tension features color-coded by fibronectin concentration.

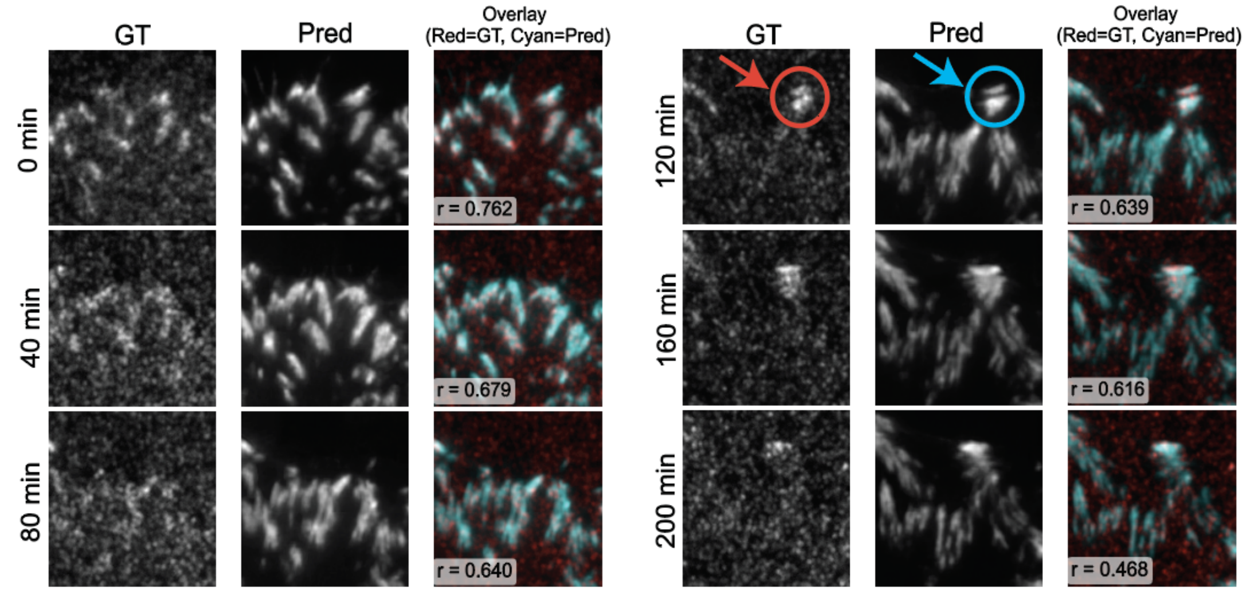


**Supplementary Figure 10: TensionDL detects emergent force hotspots on heterogeneous surfaces over long time scales.**
Ground-truth (GT) MTP images, TensionDL predictions (Pred), and GT-Pred overlays are shown for cells imaged over 200 min. Early time points (0-80 min) demonstrate strong correspondence between GT and predicted tension patterns, reflected in high GT-Pred correlations (r values shown). At later time points when many probes have photobleached (120-200 min), pearson correlations go down significantly. TensionDL nonetheless identifies newly developed force hotspots (blue arrow) and maintains sensitivity even when GT MTP signals diminish due to probe rupture or photobleaching (red arrow). At later timepoints, TensionDL predictions are likely more accurate than MTP measurements due to the effect of photobleaching and probe rupture.

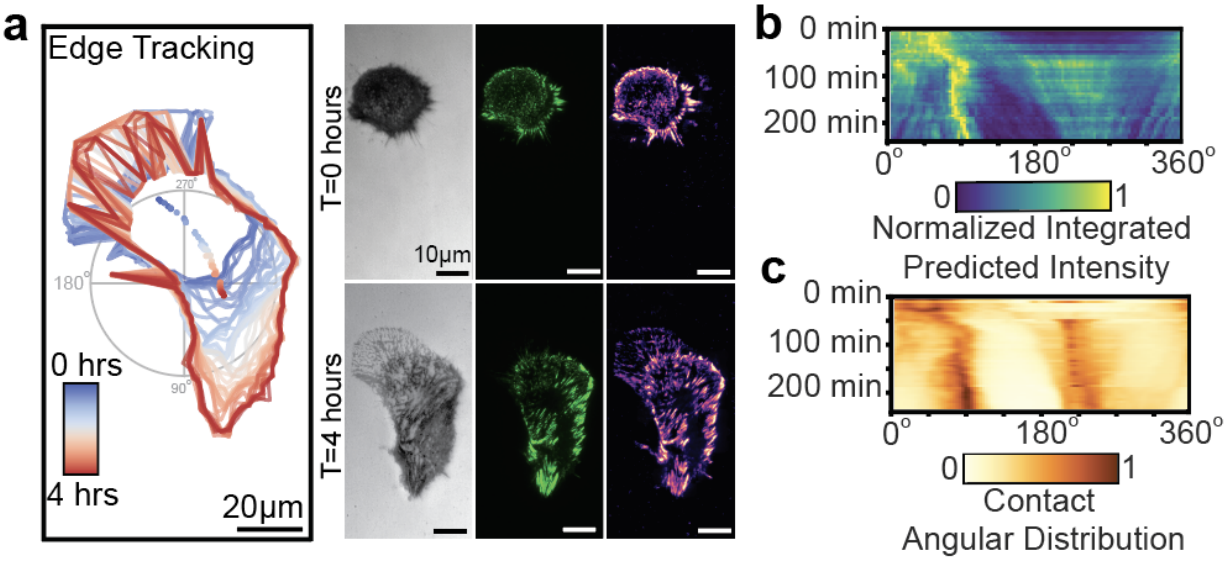


**Supplementary Figure 11: TensionDL predicts force distributions during cell migration on fibronectin-coated MTP surfaces.**

**a.** Edge tracking of the TensionDL-predicted tension map of a migrating MEF GFP–vinculin cell on fibronectin-coated MTP surfaces over 4 h; color indicates the time of observation. Right: corresponding RICM, GFP–vinculin, and TensionDL-predicted tension maps at early (t = 0 min) and late (t = 240 min) time points. Scale bar, 10 μm. **b.** Angular kymograph of normalized integrated TensionDL-predicted intensity, binned by angle (x-axis) and tracked over the 4 h migration period (y-axis). **c.** Matched angular kymograph of RICM-derived contact distribution for the same cell

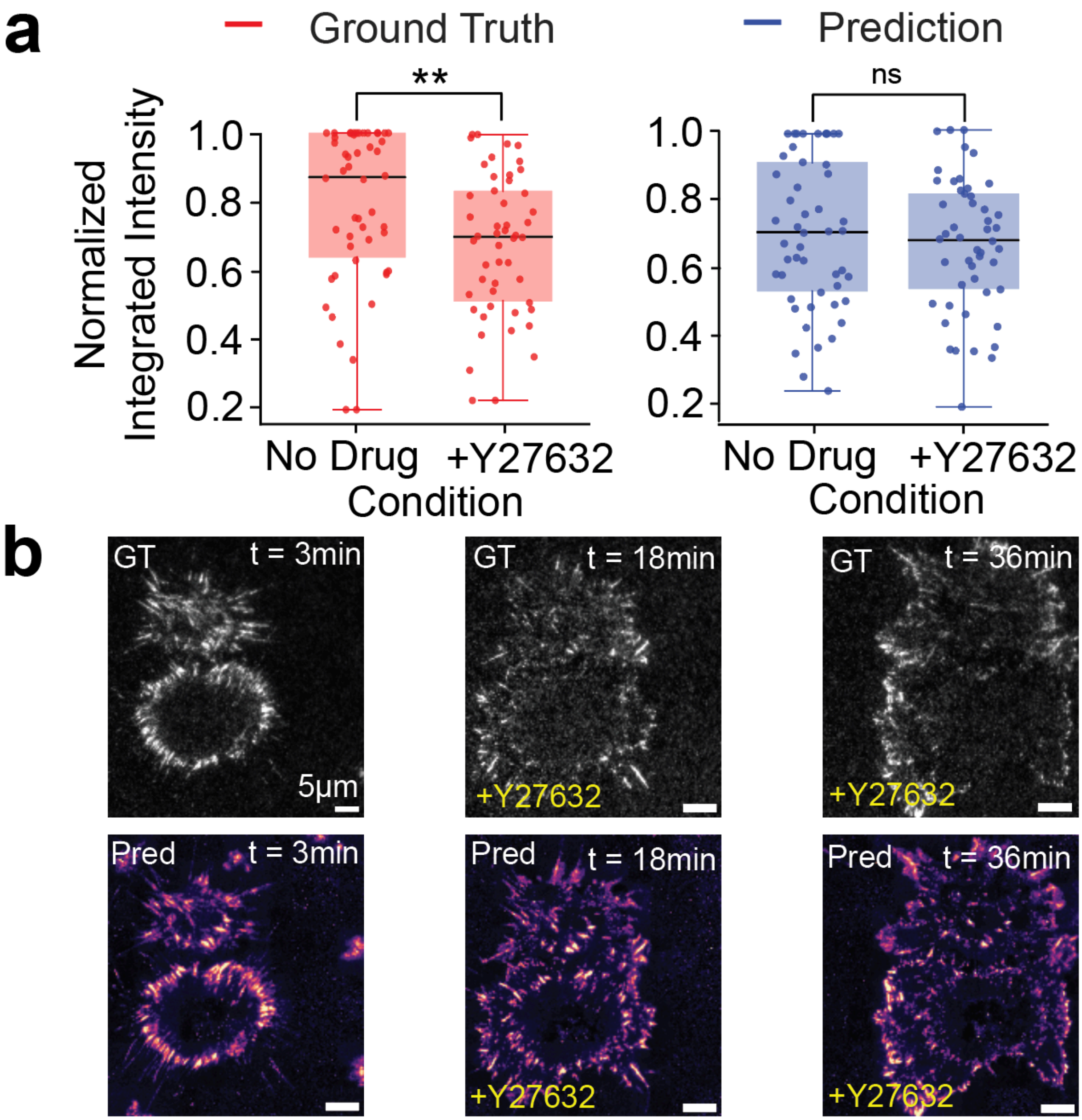


**Supplementary Figure 12: TensionDL response to Y-27632 treatment.**
**a.** Box-and-whisker plots of normalized integrated MTP-tension before and after addition of Y-27632; significance was determined by Welch's t-test (**P < 0.01; ns, not significant). **b.** Representative GT MTP-tension maps (top) and corresponding TensionDL predictions (bottom) at t = 3, 18, and 36 min. Scale bar, 5 μm.

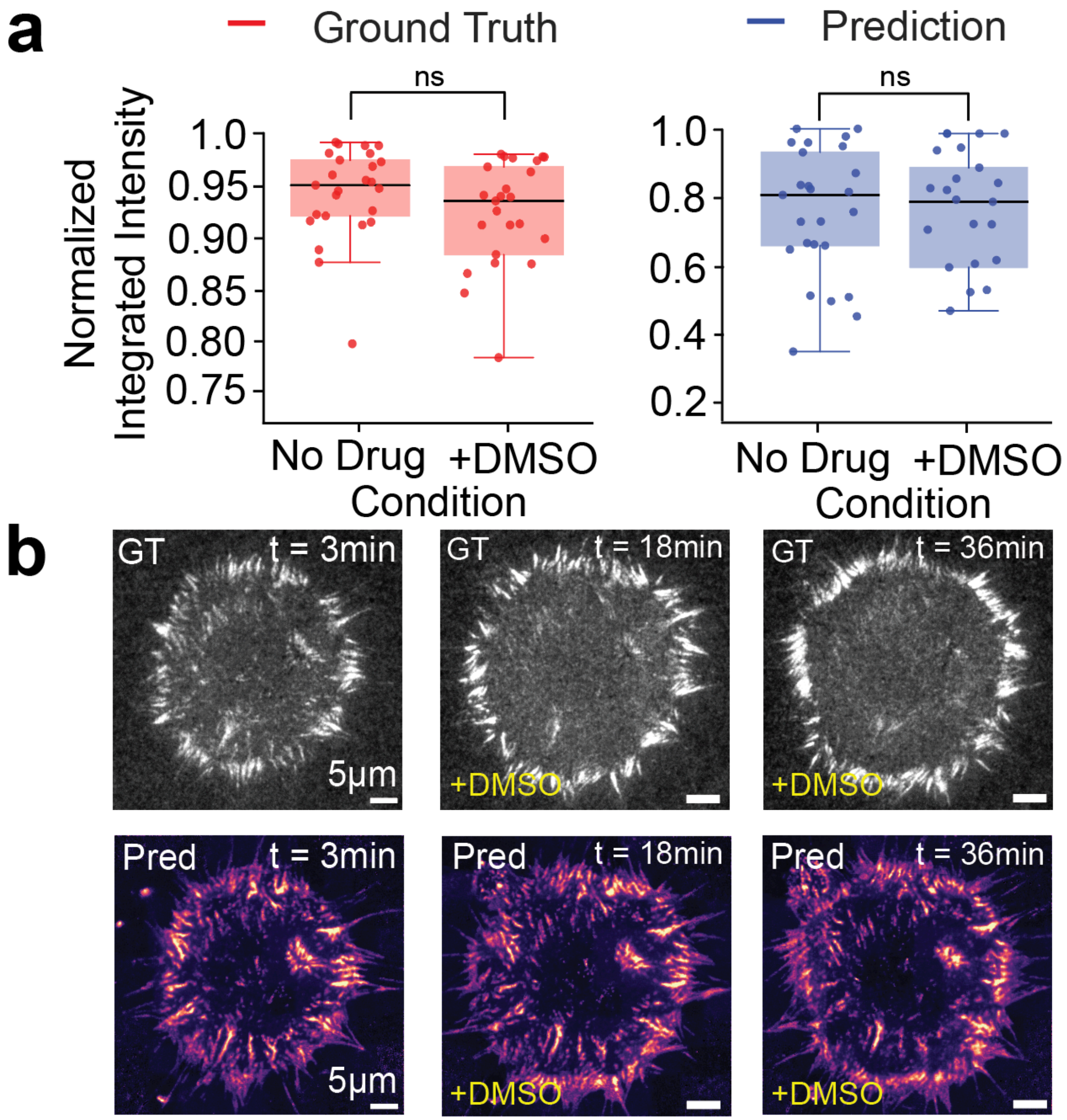


**Supplementary Figure 13: TensionDL response to Vehicle control: DMSO treatment.**
**a.** Box-and-whisker plots of normalized integrated MTP-tension before and after addition of DMSO as a vehicle control; significance was determined by Welch's t-test (ns, not significant).
**b.** Representative GT MTP-tension maps (top) and corresponding TensionDL predictions (bottom) at t = 3, 18, and 36 min. Scale bar, 5 μm.

### Supplementary Tables

**Supplementary Table 1**

| **Sequence ID** | **Sequence (5’ to 3’)** |
|---|---|
| Ligand Strand | /55OctdU/TTTGCTGGGCTACGTGGCGCTCTT/3AmMO/ |
| Anchor Strand | /5BHQ_1/CGCATCTGTGCGGTATTTCACTTT/3BioTEG/ |
| 22% GC Hairpin | GTGAAATACCGCACAGATGCGTTTGTATAAATGTTTTTTTCATTTATACTTTAAGAGCGCCACGTAGCCCAGC |
| Locking strand | GAAAAAAACATTTATACCCTACCTA |

**Supplementary Table 2**

| **Imaging Channel** | **Laser Power** | **Exposure Time** |
|---|---|---|
| 561nm | 50% | 200ms |
| 488nm | 9% | 200ms |
| RICM – 385nm | 25% | 5ms |

**Supplementary Table 3**

| **Model Version** | **Description** | **Figures** |
|---|---|---|
| TensionDL | Model trained on both Vinculin and RICM, use for qualitative predictions | Figure 1, 2, 4 |
| TensionDL-Vinc | Model trained only on Vinculin | Figure 3 |
| TensionDL-Q | Model trained on both Vinculin and RICM, use for quantitative predictions | Figure 5 |

**Supplementary Table 4**

| **Acrylamide (%)** | **Bis-acrylamide (%)** | **Acrylic acid (%)** | **Young’s modulus, E (kPa)** |
|---|---|---|---|
| 12 | 0.2 | 0.6 | 21.95 ± 1.35 |
| 12 | 0.35 | 0.6 | 35.26 ± 4.88 |
| 12 | 0.5 | 0.6 | 53.29 ± 7.26 |
